# Constitutive Expression of the Progesterone Receptor Isoforms Promotes Hormone-Dependent Development of Ovarian Neoplasms

**DOI:** 10.1101/816934

**Authors:** Margeaux Wetendorf, Rong Li, San-Pin Wu, Jian Liu, Chad J. Creighton, Tianyuan Wang, Kyathanahalli S. Janardhan, Cynthia J. Willson, Rainer B. Lanz, Bruce D. Murphy, John P. Lydon, Francesco J. DeMayo

## Abstract

Abnormal expression of the progesterone receptor (PGR) isoforms, PGRA and PGRB, is often observed in women with reproductive tract cancer. To assess the importance of the PGR isoform ratio in the maintenance of the healthy reproductive tract, mice with Cre recombinase-activated PGRA and PGRB transgenes were bred with the PGR^Cre^ mouse model to generate strains expressing either PGRA or PGRB in PGR positive tissues. The PGRB mice developed ovarian neoplasms at 23 weeks of age derived from ovarian luteal cells, while the PGRA expressing mice displayed a reduced frequency of tumor development. Transcriptomic analyses of the ovarian tumors revealed an enhanced AKT pathway signature, which is in agreement with expression changes found in human ovarian adenocarcinoma. Effective treatment with the PGR antagonist RU486 reduced tumor growth and the expression of cell cycle genes. We concluded that tumor growth and proliferation is hormone and PGR isoform dependent. Further analysis of the PGRB cistrome identified binding events of critical mitotic phase entry genes. This work suggests an intriguing mechanism whereby the expression of the PGR isoforms determines *in vivo* neoplasia through high-jacking of the cell cycle pathway.

## Introduction

Proper functioning of the reproductive tract depends on the appropriate signaling and expression of the hormone receptors and their ligands for pregnancy and the maintenance of a healthy fertile state^1^. Disruption of hormone signaling not only results in infertility and miscarriage, but also can lead to diseases such as leiomyoma, endometriosis, and reproductive tract cancer^2^. Endometrial cancer and ovarian cancer collectively contribute to over 37,000 deaths per year in the United States^3^. Further understanding of hormone signaling in the reproductive tract, especially as it relates to the initiation and progression of disease, can accelerate the development of specialized therapies to treat these diseases and cancers.

The ovary produces the female steroid hormones, estrogen and progesterone under the direction from gonadotropins secreted from the pituitary gland^1^. These hormones function by binding to their cognate receptors, the estrogen and progesterone receptors, to elicit activation or repression of their respective target genes. The progesterone receptor (PGR) is expressed in all uterine compartments, yet is limited to the pre-ovulatory granulosa cells of the ovary^4^. PGR function is further regulated through the unique expression of different isoforms^5, 6^. Genetically engineered mice have been generated with specific ablation of both isoforms: PGRAKO and PGRBKO. Phenotypic analysis of these mice demonstrated that the PGRA isoform is critical for female mouse fertility. The PGRBKO mice were fertile with normal ovulation and embryo implantation, yet displayed a mammary gland phenotype of altered ductal epithelial proliferation^7^. The only uterine phenotype attributed to the PGRB isoform in the mouse was the ability to promote epithelial proliferation in the absence of the PGRA isoform^8^. However, the reported proliferative role of progesterone within endocrine organs is variable and somewhat contradictory^2, 9, 10^.

Alterations in progesterone signaling such as irregular hormone levels, decreased receptor activity, or even abnormal isoform expression ratios can result in aberrant proliferation and reproductive-associated disease^2^. Within endometrial cancer, dominance of either PGR isoform over the other is an early biomarker for tumorigenesis^11^. Furthermore, the PGRB isoform is expressed at high levels in ovarian cancers (reviewed in^2^). Therefore, we hypothesized that alteration of PGR isoform expression would have a detrimental impact on reproductive tract homeostasis. To test this hypothesis, conditional *PgrA* and *PgrB* expression alleles were utilized as previously described^12, 13^. Mice positive for either expression allele were individually mated to *Pgr^cre^* mice^14^, resulting in the constitutive expression of either the PGRA or PGRB isoform in PGR positive tissues. PGRB expressing mice developed poorly differentiated ovarian neoplasms at 23 weeks of age that regress upon treatment with the PGR inhibitor RU486 (mifepristone).

PGRA expressing mice also develop similar ovarian neoplasms, yet at a much lower frequency. Transcriptomic profiles from the PGRB expressing ovarian neoplasms exhibit a highly proliferative signature, similar to human ovarian adenocarcinoma. These mouse models provide a new perspective regarding the function of the PGR isoforms in the initiation and progression of solid tumors in endocrine tissues.

## Methods

### Generation of the PGRB Overexpression Mouse Model

Mice were cared for according to protocol within the Institution of Animal Care and Use Committee (IACUC) at Baylor College of Medicine and the Animal Care and Use Committee at the National Institute of Environmental Health Sciences. Mice exhibiting the conditional overexpression allele for murine PGRB were described previously (*mPgrB^LsL/+^*)^13^ and crossed with *Pgr^cre/+^* mice to generate *Pgr^cre/+^mPgrB^LsL/+^* mice. *Pgr^cre/+^mPgrA^LsL/+^* mice were utilized and described previously^12^.

### Ovarian Tissue Collection

*Pgr^cre/+^* and *Pgr^cre/+^mPgrB^LsL/+^* mice at 13, 23, 28, and 33 weeks of age and *Pgr^cre/+^mPgrA^LsL/+^* mice at 33 weeks of age were euthanized at diestrous stage. The estrous stage was determined by vaginal cytology^15^. Ovarian tissue was either frozen and utilized for RNA isolation or fixed in 4% PFA at 4°C overnight and stored in 70% ethanol at 4°C for subsequent tissue processing.

### Identifying Cre Recombinase Expression Using the *Rosa^mT/mG^* Model

*Rosa^mT/mG^* mice (Jackson Laboratory) were crossed with *Pgr^cre/+^* mice to generate *Pgr^cre/+^Rosa^mT/mG^* mice. Female *Pgr^cre/+^Rosa^mT/mG^* mice were euthanized during diestrus, as established by vaginal cytology. The ovaries were embedded in the optimal cutting temperature (O.C.T.) compound (Tissue-Tek, VWR 102094-104) and immediately frozen on dry ice. Frozen sections (10µm) were sectioned using the cryostat (Leica, Germany), then incubated at 65°C for 10 minutes. The endogenous Tdtomato and GFP fluorescence were visualized by Epifluorescence microscopy (Zeiss, USA).

### Ultrasound Imaging and Tumor Volume

Ultrasound was performed using the VisualSonics 2100 System at the Mouse Phenotyping Core located at Baylor College of Medicine to identify the presence of abnormal masses within the reproductive tract. For each mass, images were processed according to VisualSonics software instructions to determine tumor volumes. Tumor volume was calculated via serial contouring of the tumor cross sections according to the VisualSonics 2100 3D tumor imaging software.

### RU486 Inhibition of Tumor Growth

After identification of tumor presence using ultrasound imaging, mice exhibiting a tumor with at least a 50mm^3^ tumor volume were administered a chronic release pellet of RU486 (30mg/pellet, 60-day release) or placebo pellet from Innovative Research of America by subcutaneous placement at the nape. The mice were imaged weekly for 9 weeks via ultrasound 3D video imaging to track the tumor size. ImageJ was used to assess area, cell signal, and integrative density measurements for the middle region of each tumor at the final week of treatment (measurement analysis on ImageJ 1.49 software, U.S. National Institute of Health, Bethesda, MD, USA). Mice were sacrificed 8 weeks after pellet surgery or until the tumor size reached 10% of the mouse body weight. Fold tumor volume change for vehicle and RU486 treated tumors was calculated as the average fold change (log10(tumor volume) normalized to one) for each treatment.

### Histology and Immunohistochemistry

Tissues were fixed in 4% PFA and embedded in paraffin wax. Embedded tissues were sectioned at 5µm and incubated for 20 minutes at 60°C. After 10 minutes of cooling, the slides were dewaxed in xylenes and a decreasing gradient of pure ethanol. For hematoxylin and eosin (H&E) staining, tissues were adequately stained with H&E and were then dehydrated before coverslips were applied. Ovarian histopathology of H&E stained sections was evaluated by two board-certified veterinary pathologists.

For immunohistochemistry, antigen retrieval was performed according to manufacturer’s instructions (Vector Labs Antigen Unmasking Solution H-3300). Endogenous peroxide was blocked using 3% hydrogen peroxide diluted in methanol. The tissue was blocked before application of primary antibody overnight (PGR: Dako A0098 (1:1000 Supplemental Figure 1, 1:400 Figure 2), Myc-tag: Cell Signal 71D10 (1:150), BrdU: BD Pharmingen Cat# 551321 (1:50), AKT: Cell Signal 4691 (1:400), pAKT: Cell Signal 4060 (1:100), Ki67: Abcam ab15580 (1:1000), cleaved CASP3: Cell Signal 9661 (1:400), CCND1: NeoMarkers RB-9041-P0 (1:400)). Secondary antibody was diluted in 1% BSA at a concentration of 1:200. The ABC reagent was applied to tissues according to manufacturer’s instructions (Vector Labs ABC PK- 6100). Signal was developed using Vector Labs DAB Immpact staining according to manufacturer’s instructions (Vector Labs SK-4105). Tissue was counterstained with hematoxylin and dehydrated before applying coverslips.

**Figure 1:**
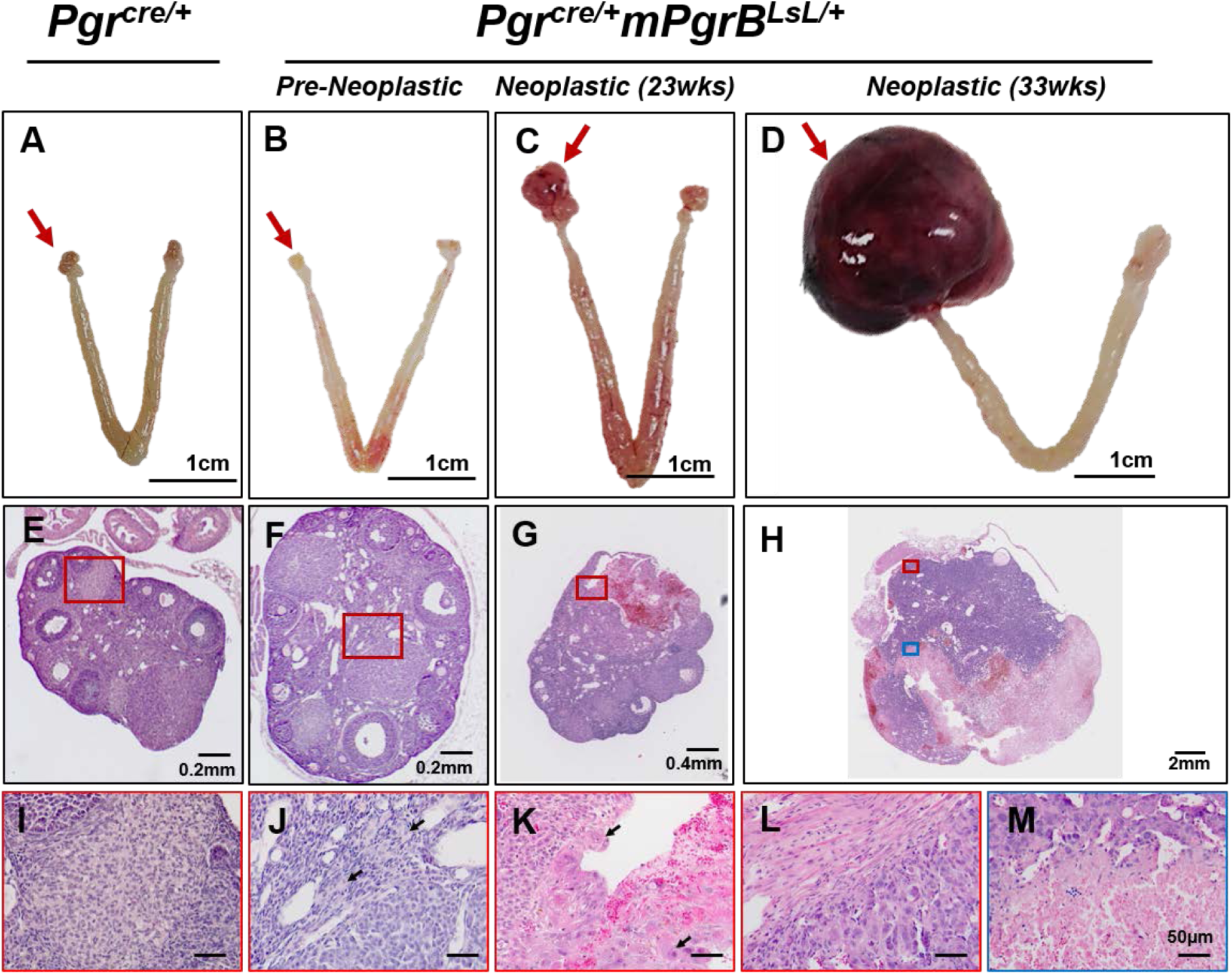
Constitutive PGRB expression induces ovarian tumors in mice. Pictures of ovary in *Pgr^cre/+^* mice (A) and pre-neoplastic and neoplastic at 23 and 33 weeks of age in ovarian tumors from *Pgr^cre/+^mPgrB^LsL/+^* mice (B,C,D). Hematoxylin and eosin stained sections of ovaries in *Pgr^cre/+^* mice (E) and pre-neoplastic and neoplastic at 23 and 33 weeks of age from *Pgr^cre/+^mPgrB^LsL/+^* mice (F,G,H). Higher magnifications of selected regions in the histology of ovaries in *Pgr^cre/+^* mice (I) and pre-neoplastic and neoplastic at 23 and 33 weeks of age from *Pgr^cre/+^mPgrB^LsL/+^* mice (J,K,L,M). Black arrows indicate tumor cells with big nuclear size and/or big cell size. Red arrows indicate the imaged ovary or ovarian tumor.

**Figure 2:**
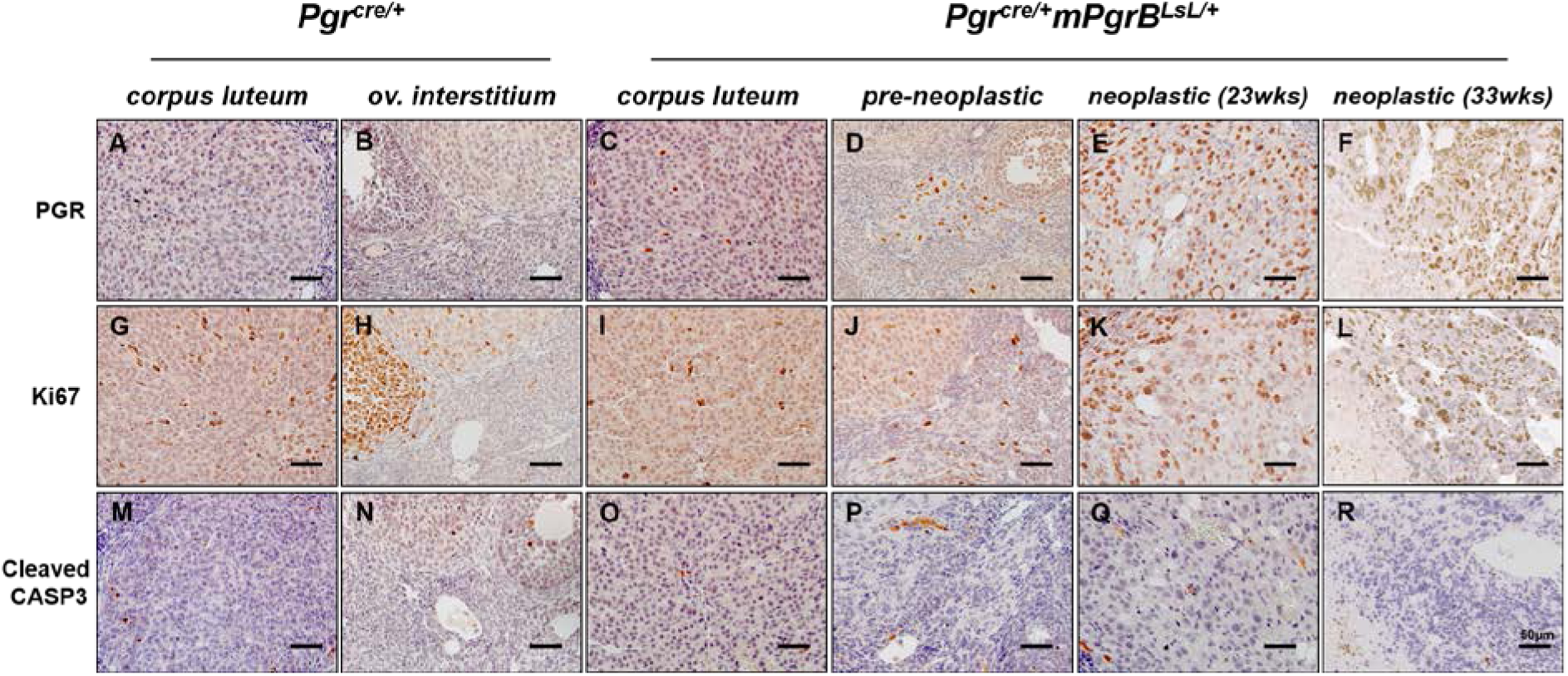
PGR and Ki67 positive staining in ovarian neoplasia from *Pgr^cre/+^mPgrB^LsL/+^* mice. Immunohistochemistry of PGR (A-F), Ki67 (G-L), and cleaved caspase 3 (CASP3) (M-R) in *Pgr^cre/+^* mouse corpus luteum (A,G,M) and ovarian interstitial tissue (B,H,N) and *Pgr^cre/+^mPgrB^LsL/+^* mouse normal corpus luteum (C,I,O), pre-neoplastic abnormal cells (D,J,P), neoplastic tumor cells from 23 weeks of age (E,K,Q), and neoplastic tumor cells from 33 weeks of age (F,L,R) using DAB chromogen with hematoxylin counterstain.

### TUNEL Staining

Tissues were fixed in 4% PFA and embedded in paraffin wax. Embedded tissues were sectioned at 5µm and incubated for 20 mins at 60°C. After 10 minutes of cooling, the slides were dewaxed in xylenes and a decreasing gradient of pure ethanol. Tissues were incubated at 37°C for 1 hour with a 20mg/mL proteinase K solution diluted in 10mM Tris/HCl pH 7.4-8. Tissues were washed in PBS and labeled using the Roche In Situ Cell Death TMR Red kit (Roche Diagnostics) according to manufacturer’s instructions. Slides were cover-slipped with Vectashield+DAPI (Vector Labs) and sealed with clear nail polish.

### BrdU and TUNEL Quantification

BrdU labeling reagent was i.p. injected two hours before euthanasia (GE Healthcare). Regions of the 40x (BrdU) and 20x (TUNEL) images were marked and quantified for total cells and total positive DAB stained cells (BrdU) or total positive rhodamine red puncta (TUNEL). Percent positive cells were quantified for each sample and averages were quantified to represent the mean percent positive cells for BrdU nuclei or TUNEL positive puncta.

### Western blots

Isolated protein was applied to a Bis-Tris NuPAGE 4%-12% gel (Novex by Life Technologies) for protein separation. Protein was wet transferred to a polyvinylidene difluoride membrane and blocked in 5% blotting grade nonfat milk diluted in phosphate-buffered saline (PBS) with 0.1% Tween for at least one hour. Membranes were incubated with primary antibody (PGR Santa Cruz H-190 (1:400), Actin Santa Cruz I-19 (1:10,000), myc-tag Origene TA100010 (1:1000), Cyclin D1 Thermo Fisher RB-9041-P0 (1:200)) overnight. Membranes were washed and incubated with secondary antibody (anti-rabbit peroxidase (1:4000), anti-mouse peroxidase (1:5000), and anti-goat peroxidase (1:4000) according to primary antibody requirements) in 5% milk diluted in PBS with Tween. The Amersham ECL Western blotting system was utilized to develop peroxidase labeled protein according to manufacturer’s instructions (GE Healthcare).

### RNA Isolation

Frozen tissue was homogenized in TRIzol reagent (Thermo Fisher). RNA was isolated using chloroform and precipitated using isopropanol with resuspension in water. For RNA prepared for microarray, TRIzol reagent was utilized followed by the aqueous phase isolation using 1-Bromo-3-chloropropane and a second aqueous phase isolation using chloroform. The aqueous layer was then mixed with 100% ethanol and applied to the column from the Qiagen RNEasy RNA mini prep kit. The column was washed, and RNA was isolated using manufacturer’s instructions (Qiagen).

### Reverse Transcriptase PCR and Quantitative Real Time PCR

RNA was reverse transcribed into cDNA using M-MLV reverse transcriptase (Thermo Fisher) according to manufacturer’s instructions. Quantitative real time PCR was performed using Taqman Master Mix (Life Technologies) or SYBR Green Master Mix (Roche Diagnostics). Taqman primers and probes were acquired from Life Technologies and SYBR primers were designed based on Primer Bank predictions using Sigma-Aldrich synthesized oligonucleotides (listed in Supplemental Table 4). Delta Ct values were calculated using 18S control amplification to acquire relative mRNA levels per sample.

**Figure 4.**
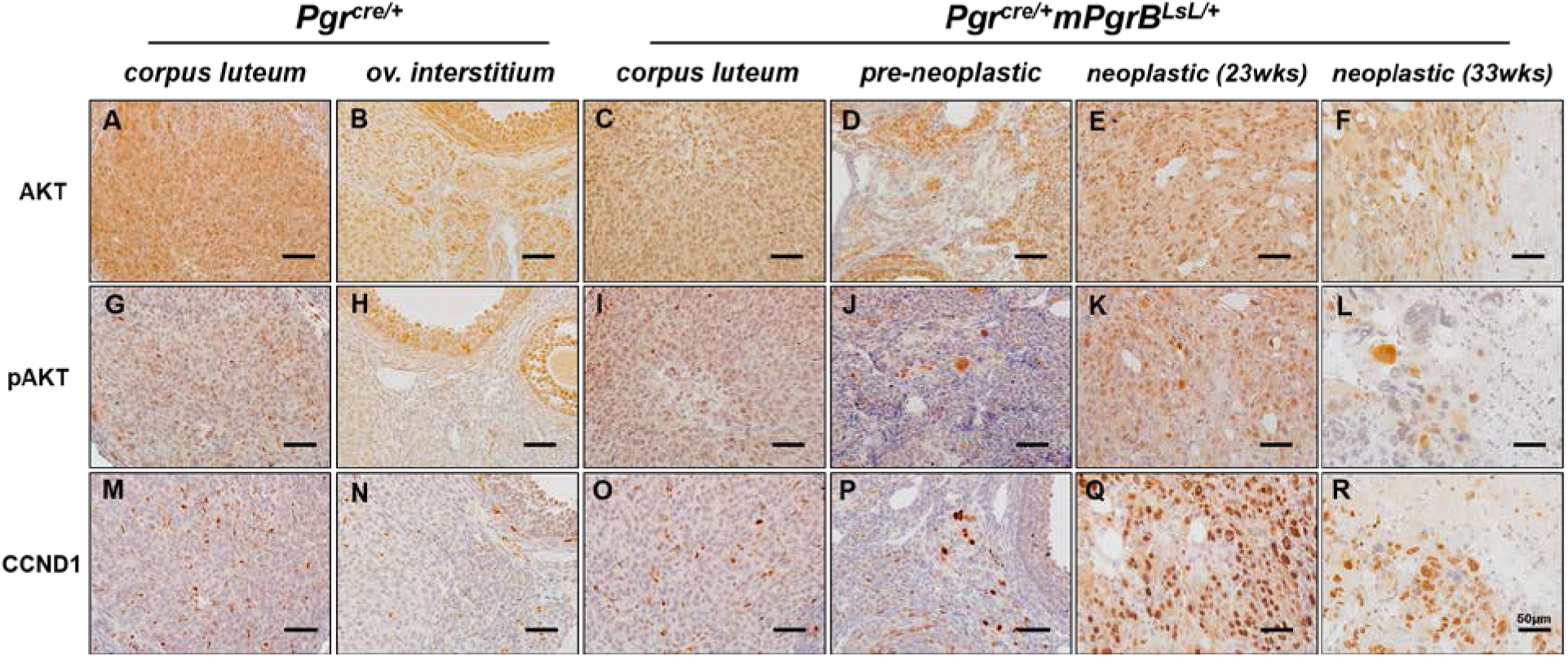
AKT, pAKT and CCND1 positive cells in ovarian neoplasia from *Pgr^cre/+^mPgrB^LsL/+^* mice. Immunohistochemistry of AKT (A-F), phosphorylated AKT (G-L), and CCND1 (M-R) in *Pgr^cre/+^* mouse corpus luteum (A,G,M) and ovarian interstitial tissue (B,H,N) and *Pgr^cre/+^mPgrB^LsL/+^* mouse normal corpus luteum (C,I,O), pre-neoplastic abnormal cells (D,J,P), neoplastic tumor cells from 23 weeks of age (E,K,Q), and neoplastic tumor cells from 33 weeks of age (F,L,R) with DAB chromogen and hematoxylin counterstain.

### RNA Microarray

RNA quality was assessed using the Agilent 2100 Bioanalyzer (Agilent Technologies). Microarrays were performed by the Genomic and RNA Profiling Core at Baylor College of Medicine and the Epigenomic Core Laboratory at the National Institute of Environmental Health Sciences. For microarrays, sample libraries were amplified and labeled, and individual cDNA samples were hybridized to either the Agilent G3 Mouse GE 8×60k array or the Affymetrix GeneChip Mouse Genome 430 2.0 array according to manufacturer’s instructions (Agilent, Affymetrix). Array data was analyzed using Bioconductor for quantile normalization. Significantly regulated genes were identified using a p-value ≤ 0.05 with a variable fold change region, identified specifically in each figure legend. All raw microarray data are available on NCBI-GEO database with accession number GSE137433.

### Chromatin Immunoprecipitation and Sequencing

Tissue was flash frozen and sent to Active Motif for Factor Path chromatin immunoprecipitation and sequencing analysis. Tissue was fixed, then sheared into small fragments before immunoprecipitation with the Active Motif PGR antibody. Bound DNA was isolated to generate a purified and amplified library of PGR bound sequence regions and sequenced using the Illumina sequencing platform. PGR bound intervals were identified using MACS analysis and mapped to genes by the Active Motif Company. The Active Motif Company also performed validation of the PGR binding events using ChIP-qPCR through measurement of the amount of binding events per cell compared to binding events occurring in an untranslated region. ChIP-Seq data is available on the NCBI-GEO database with accession number GSE137433.

### Data Analysis

Pathway analysis performed on microarray data was analyzed using Ingenuity Pathway Analysis software and the public Database for Annotation, Visualization, and Integrated Discovery (DAVID) with default settings applied. ChIP-Seq data quality, binding enrichment, and motif analysis was assessed using the Cistrome Analysis Pipeline software (http://cistrome.org/ap/). GraphPad Prism software was utilized to perform one-way ANOVA, multiple comparison test, and Student’s t-Test analyses for qRT-PCR, TUNEL quantification, and BrdU quantification data. Hormone response elements were identified using HOMER de novo motif analysis (http://homer.salk.edu/homer/). Hierarchal clustering heatmaps were generated using Partek Genomics Suite 6.6 software. NextBio (Illumina) analysis was performed on differentially regulated genes using a p-value ≤ 0.05 and an absolute fold change of >1.3 before comparison to other published studies. For Gene Set Enrichment Analysis (GSEA) at the Broad Institute MSigDB, all genes were ranked based on fold changes and analyses were performed against C6 (oncogenic signatures). Top significantly enriched gene sets were selected on the basis of false discovery rate (FDR) q-value < 0.05.

## Results

### Generation of the PGR Expression Models

To assess the importance of the PGR isoform ratio in the maintenance of the reproductive tract, PGR overexpression models were generated. The altered PGR isoform ratio was achieved by mating *mPgrA^LsL/+^*^12^ mice or *mPgrB^LsL/+^*^13^ mice to the *Pgr^cre^* mice^14^, resulting in Cre recombinase expression in PGR expressing cells. As a result, the PGRA or PGRB isoform was constitutively expressed in all compartments of the uterus compared to wildtype, in which PGR is expressed at differing levels across estrous stages and pregnancy (PGRA published in^12^, PGRB in Supplemental Figure 1A-B). Furthermore, PGRB expression was observed in scattered cells within the ovarian corpora lutea of the *Pgr^cre/+^mPgrB^LsL/+^* mice detected by myc-tag immunohistochemistry (Supplemental Figure 1C-F). To assess the total level of PGR protein within the uterus relative to the PGRB transgene expression, protein isolated from whole uterine tissue was measured for PGR and myc-tag epitope levels. Whole uterine protein isolates from *Pgr^cre/+^mPgrB^LsL/+^* mice displayed elevated levels of myc-tagged PGRB protein (118 kDa) with markedly low levels of PGRA protein (90 kDa) compared to wildtype *Pgr^cre/+^* mice (Supplemental Figure 1G).

### PGR expressing mice develop ovarian neoplasia

Surprisingly, *Pgr^cre/+^mPgrB^LsL/+^* female mice developed ovarian tumors with 100% penetrance by 23 weeks of age. *Pgr^cre/+^mPgrA^LsL/+^*also developed tumors but at a much reduced frequency of 18.2% (Table 1). A time course of tumor development was assessed by sacrificing mice at 13, 23, 28, and 33 weeks of age. Ovarian tumor development was divided into two categories. The presence of abnormal cells in the ovaries was categorized as “pre-neoplastic” while the presence of tumors was called “neoplastic”. At 23 weeks of age, the *Pgr^cre/+^mPgrB^LsL/+^* mice exhibited the presence of abnormal cells (pre-neoplastic) or visible tumors (neoplastic) in the ovarian interstitium (Table 1). The gross morphology and histological features of the ovarian pathology is shown in Figure 1. As tumor development progressed, the tumor tissue effaced the normal ovarian architecture and filled the ovarian bursa. In the initial stages, numerous large pleomorphic cells were present in and around the corpora lutea. At a later stage when the tumor is completely formed, they were composed of polygonal cells admixed with large blood-filled spaces and interstitial cells. The neoplastic cells had variably distinct borders, scant to abundant eosinophilic cytoplasm and variably sized nucleus with coarsely stippled chromatin. Marked anisocytosis and anisokaryosis was present. There were multiple, prominent nucleoli in some of the cells. Occasional multinucleated cells were present. Mitotic figures were rare. Many of the cells appeared to have a large vacuole in their cytoplasm. Based on the histomorphology, the tumors were diagnosed as poorly differentiated neoplasms of the ovary. Ovarian neoplastic cells exhibited intense staining for PGR and the proliferative marker, Ki67, in neoplastic tissues compared to wildtype ovarian corpus luteum and interstitium (Figure 2). Furthermore, the expression of the apoptotic marker, cleaved caspase 3 (CASP3), is reduced in *Pgr^cre/+^mPgrB^LsL/+^* ovarian tumor tissue. Upon further immunohistochemical analysis, PGR positive cells from regressing corpora lutea were found maintained in the ovarian stroma (Supplemental Figure 2A-D). These ovarian tumors also demonstrated high expression levels of the steroidogenic acute regulatory protein (StAR), a regulator of steroidogenesis and reminiscent of normal corpora lutea (Supplemental Figure 2E-F). To assess whether this ovarian pathology was unique to the PGRB isoform, *Pgr^cre/+^mPgrA^LsL/+^* mice were aged and assessed for the presence of tumor formation. As shown in Table 1, the *Pgr^cre/+^mPgrA^LsL/+^* mice exhibited a similar formation of ovarian pathology, yet with a much lower frequency. However, both mouse models did not exhibit any metastases. We concluded that abnormal increased expression of the individual PGR isoforms can result in ovarian neoplasia with increased proliferation and decreased apoptosis, potentially derived from corpora luteal cells. We next wanted to further understand the cell type-specific origin of the ovarian PGR isoform expression.

**Table 1:**
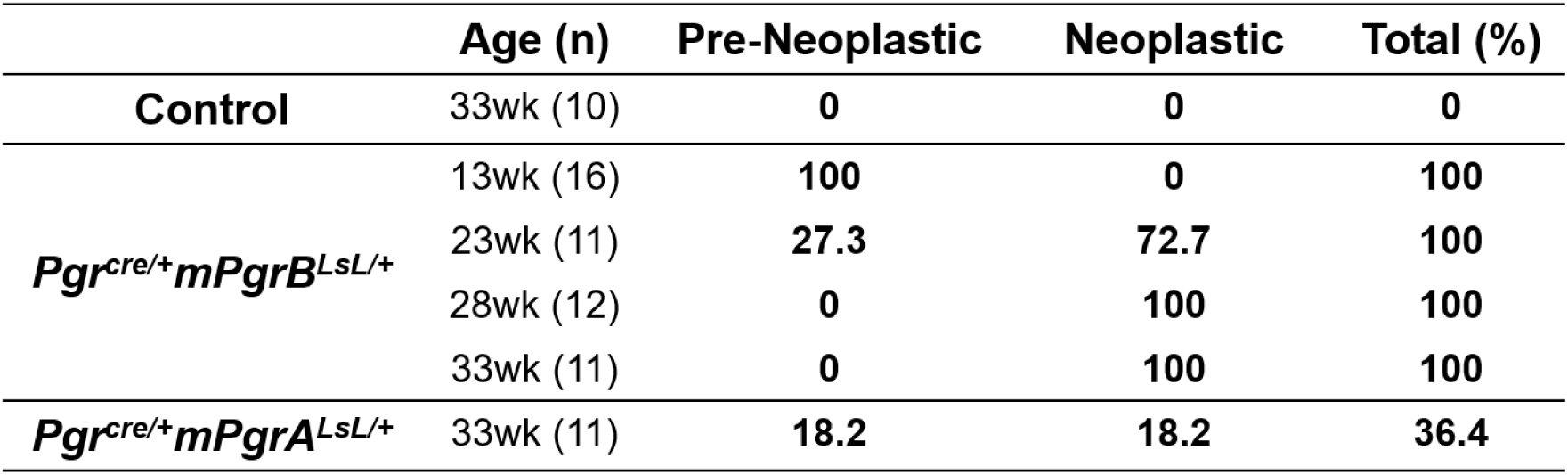
Time Course of Ovarian Tumor Development in Pgr^cre/+^mPgrA^LsL/+^ mice and Pgr^cre/+^mPgrB^LsL/+^ mice. wk: weeks. Number in parenthesis: number of total mice.

### *Pgr^cre/+^Rosa^mT/mG^* mouse model demonstrates the presence of Cre recombination in regressing corpora lutea

Endogenous PGR is expressed in the granulosa cells of the pre-ovulatory follicle. To determine the cell lineage depicting *Pgr^Cre^* activity in the mouse ovary, *Pgr^cre/+^Rosa^mT/mG^* mice were implemented. In this model, expression of the mTomato (red) marker indicates cells with no Cre recombinase activity. Cells expressing GFP (green) demarcates the cells in the lineage of *Pgr^cre^* expression.^16^. To identify where the *Pgr^cre/+^* initiates recombination and resultant PGRB expression in the ovary, fluorescence was measured in ovarian tissue from *Pgr^cre/+^mPgrB^LsL/+^* mice. Positive green staining demonstrates recombination occurring in regressing corpora lutea of the ovary (Supplemental Figure 3). Thus, the origin of the *Pgr^cre/+^mPgrB^LsL/+^* mouse ovarian neoplasms is likely within the regressing corpora lutea at the site of recombination.

**Figure 3:**
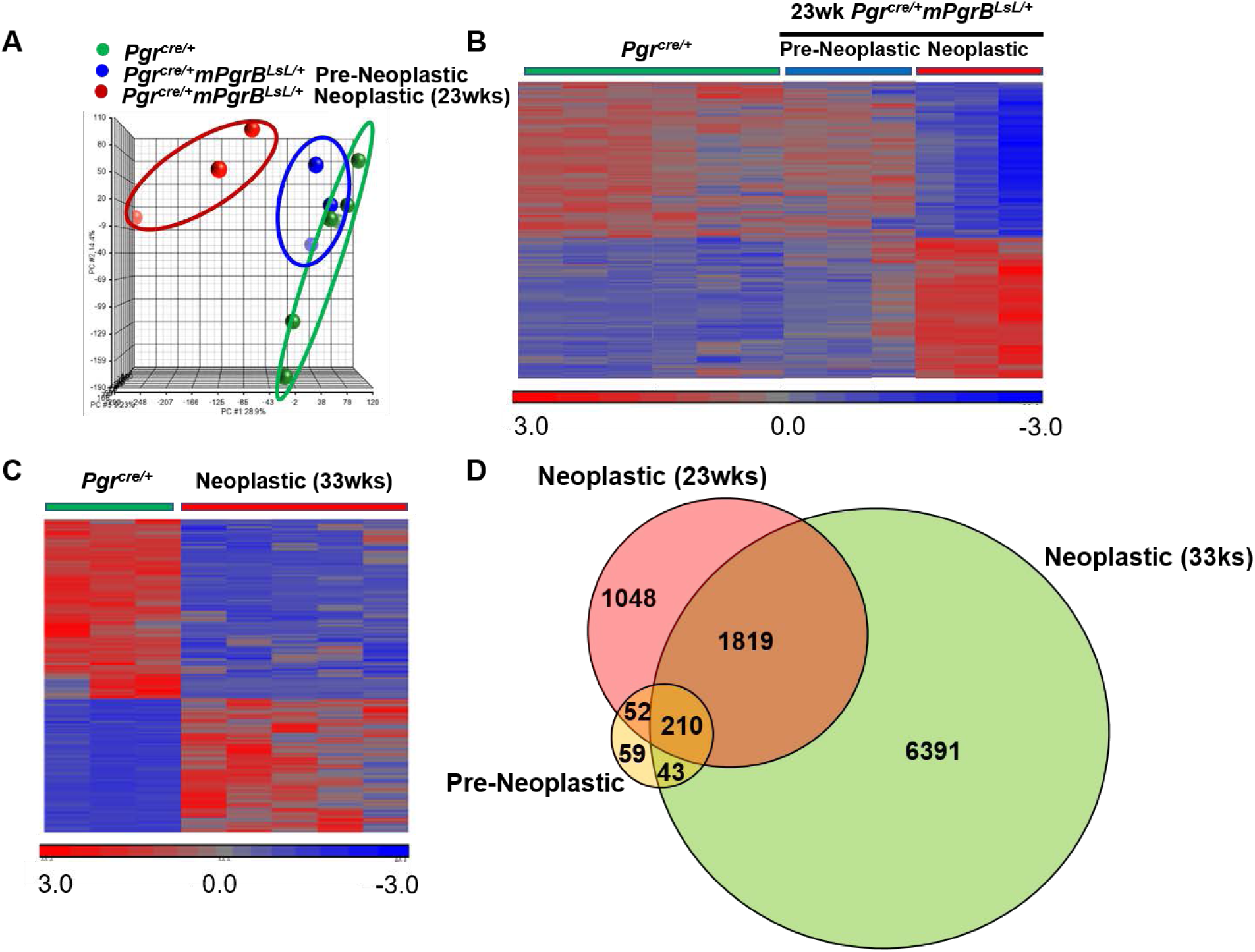
Gene transcriptome alterations in *Pgr^cre/+^mPgrB^LsL/+^* mice during tumor progression. (A) Principle component analysis (PCA) identified two clusters in 23 week old *Pgr^cre/+^mPgrB^LsL/+^* mouse ovaries, defined as pre-neoplastic and neoplastic. The pre-neoplastic cluster exhibited similar characteristics to *Pgr^cre/+^* mouse ovaries. However, the cluster from neoplastic, 23 week old *Pgr^cre/+^mPgrB^LsL/+^* mouse ovarian tumor tissue exhibited a unique signature compared to *Pgr^cre/+^* mouse ovaries. (B) Transcriptome heatmap demonstrating shifts in expression levels from pre-neoplastic to neoplastic ovarian tissue from 23 week old *Pgr^cre/+^mPgrB^LsL/+^* mice. (C) Transcriptome heatmap of neoplastic ovarian tissue from 33 week old *Pgr^cre/+^mPgrB^LsL/+^* mice compared to wildtype *Pgr^cre/+^* mouse ovaries. (D) Venn diagram depicting differentially regulated genes in pre-neoplastic, neoplastic at 23 weeks of age, and neoplastic at 33 weeks of age from *Pgr^cre/+^mPgrB^LsL/+^* mouse ovarian tissue with a p-value < 0.05, fold changes <-1.3 and >1.3.

### Cycling progesterone levels and FSH and LH levels are normal in conditional PGR expression mice

Although the PGR is genetically engineered to express at high levels in the *Pgr^cre/+^mPgrB^LsL/+^* mice and the previously described *Pgr^cre/+^mPgrA^LsL/+^* mice^12^, the effect of the conditional allele on circulating progesterone ligand is unknown. Excess levels of cycling progesterone in mice can dramatically affect the amount of active PGR signaling. To identify the levels of endogenous progesterone levels, serum was obtained from virgin, 23-week old *Pgr^cre/+^mPgrB^LsL/+^* and *Pgr^cre/+^mPgrA^LsL/+^* mice and submitted to the Ligand Assay and Analysis Core at the Center for Research in Reproduction at the University of Virginia. The level of cycling progesterone across all mice was comparable with no significant differences (Supplemental Figure 4A). Additionally, follicle-stimulating hormone (FSH) and luteinizing hormone (LH), which signal from the pituitary to the ovary to produce the steroid hormones^1^, also exhibited normal levels across all genotypes (Supplemental Figure 4B-C). Thus, the PGRB and PGRA overexpression alleles do not impair circulating progesterone ligand or FSH and LH levels.

### Comparative Transcriptomic Analysis of Tumor Development Stages from *Pgr^cre/+^mPgrB^LsL/+^* Mice

To identify changes in gene expression during tumor progression, ovarian tumors were harvested from *Pgr^cre/+^mPgrB^LsL/+^* mice, while age-matched ovaries from wildtype *Pgr^cre/+^* mice served as control. Total RNA was isolated and RNA microarrays were performed. Principle component analysis on the raw microarray data for wildtype, pre-neoplastic, and neoplastic stages at 23 weeks of age (Figure 3A) revealed the transcriptome from *Pgr^cre/+^* tissue was similar to the profile from pre-neoplastic tissue samples. However, the neoplastic transcriptome profile exhibited an overall differential expression compared to wildtype or pre-neoplastic profiles. These same results were confirmed in RNA profiling heatmap analysis depicted in Figure 3B. Interestingly, the pre-neoplastic expression profile, although similar to the wildtype profile, also exhibits some similarity to the neoplastic profile (Figure 3B). Furthermore, separate transcriptomic analyses between 33 week old *Pgr^cre/+^* ovarian tissue and neoplastic tumor tissue demonstrate a profile very similar to the changes observed between 23 week old wildtype and neoplastic profiles (Figure 3C, 3B). Therefore, the 23 and 33 week old neoplastic tumor RNA expression profiles differ from age-matched wildtype ovarian tissue. To further compare these microarray profiles, genes with a fold change of greater than 1.3 and less than −1.3 with a p-value ≤ 0.05 from each tumor stage expression profile were intersected and are shown in Figure 3D in Venn diagram format. Neoplastic profiles from both 23 and 33 weeks of age exhibit strong similarity with 2,029 genes overlapped, with 210 of those genes also observed in the pre- neoplastic expression profile. Total lists of differentially expressed genes for pre-neoplastic, neoplastic at 23 weeks of age, and neoplastic at 33 weeks of age are reported in the supplemental Excel spreadsheet, workbooks 1.1-1.3. Further examination was performed using Ingenuity Pathway Analysis to identify the top altered canonical pathways in neoplastic ovarian tissue at 23 weeks of age. This expression profile exhibits strong characteristics for proliferation, growth, and immune cell signaling (Supplemental Table 1). Notably, the PI3K/AKT pathway and cell cycle pathway were significantly changed, representing two routes of regulation of cell growth^17^. Thus, at 23 weeks of age, the neoplastic ovarian tumor tissue exhibits a strong growth profile, indicative of neoplasia.

### AKT signaling is upregulated in ovarian tissue from *Pgr^cre/+^mPgrB^LsL/+^* mice

Due to the strong PI3K/AKT signature identified in the top canonical pathway list for neoplastic ovarian tissue at 23 weeks of age (Supplemental Table 1), immunohistochemistry on all stages of ovarian tissue from *Pgr^cre/+^mPgrB^LsL/+^* mice was examined for active AKT signaling. Staining for AKT and active, phosphorylated AKT was performed on wildtype ovarian corpus luteum and interstitium and compared to corpus luteum from pre-neoplastic and both neoplastic stages ovarian interstitium from *Pgr^cre/+^mPgrB^LsL/+^* mice. AKT and pAKT positive staining was observed in neoplastic tissue from *Pgr^cre/+^mPgrB^LsL/+^* mice (Figure 4A-L). Additionally, CCND1, a critical cyclin protein required for S phase entry into the cell cycle, was identified to be dramatically upregulated in neoplastic ovarian tissue at both 23 and 33 weeks of age from *Pgr^cre/+^mPgrB^LsL/+^* mice compared to wildtype (Figure 4M-R). Therefore, PGRB expression in the ovary results in heightened levels of active AKT and CCND1, critical regulators of cell proliferation pathways.

To further examine the similarity between the ovarian tissue from *Pgr^cre/+^mPgrB^LsL/+^* mice and that from ovarian cancer patients, Gene Set Enrichment Analysis (GSEA) was performed on the gene expression profile from 33 week old neoplastic ovarian tissue. Positive enrichment for the AKT pathway was identified in the top 2 enriched gene lists in common between human ovarian adenocarcinoma and neoplastic ovarian tissue (Figure 5A). Additionally, NextBio gene expression profile analysis identified more than half of the significantly changed genes were similar to published human ovarian adenocarcinoma datasets (Figure 5B). Out of these 4,211 genes shared between the human dataset and the advanced neoplastic array, the majority of the genes were identified to be changed in the same direction (Figure 5C). Thus, differential expression changes identified from advanced neoplastic tissues harvested from *Pgr^cre/+^mPgrB^LsL/+^* mice exhibit a similar transcriptomic profile compared to human ovarian adenocarcinoma.

**Figure 5:**
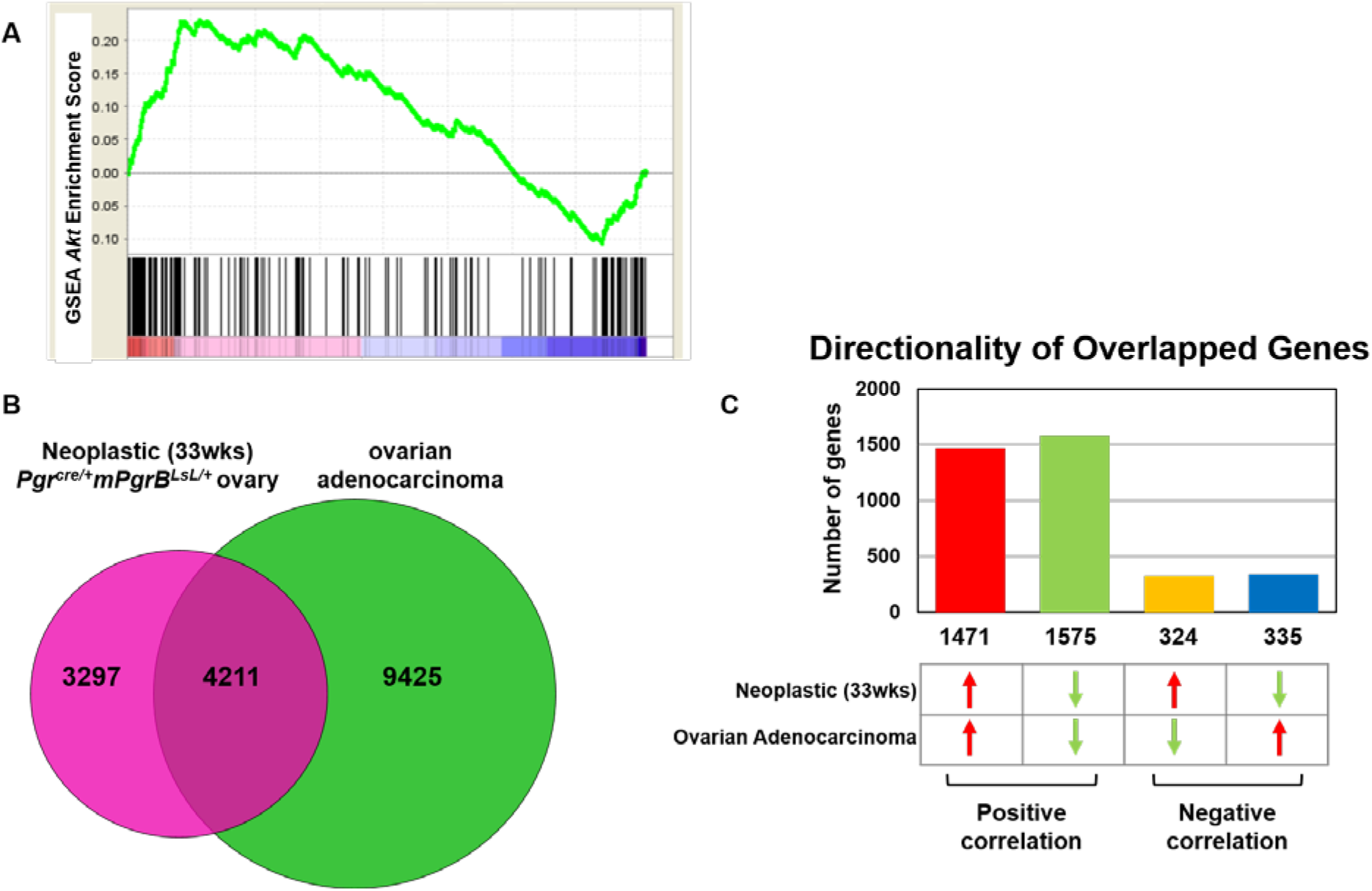
PTEN/Akt signaling pathway is highly enriched in neoplastic *Pgr^cre/+^mPgrB^LsL/+^* mouse ovarian tissue transcriptome data. (A) Gene Set Enrichment Analysis (GSEA) shows Protein Kinase B (AKT) signaling is highly enriched in *Pgr^cre/+^mPgrB^LsL/+^* mouse neoplastic ovarian tissue at 33 weeks of age. (B-C) NextBio analysis on transcriptome from neoplastic *Pgr^cre/+^mPgrB^LsL/+^* mouse ovarian tissue at 33 weeks of age identifies overlap with a murine ovarian adenocarcinoma dataset. Utilizing NextBio analysis, the 33 week old, neoplastic *Pgr^cre/+^mPgrB^LsL/+^* mouse ovarian tissue transcriptome data exhibited similarity to an ovarian adenocarcinoma microarray dataset resulting from deletion of PTEN and APC in the ovarian epithelium cited in Wu., R *et al.*, *Cancer Cell 11*, 321-333, April 2007. (B) Venn diagram depicting overlap of significantly changed genes in 33 week old, neoplastic *Pgr^cre/+^mPgrB^LsL/+^* mouse ovarian tissue transcriptome data in pink compared to the murine ovarian adenocarcinoma microarray by Wu R., *et al* in green with an overlap p-value of 1.4E-65. (C) Description of gene number and direction (upregulated or downregulated) for the 4211 overlapped genes in (B).

### Chronic RU486 treatment results in the regression of tumor volume in PGRB expressing mice

Due to the robust phenotype of abnormal cells and developing tumors in the *Pgr^cre/+^mPgrB^LsL/+^* mice, it was important to determine whether progesterone signaling was responsible for driving tumor growth in these mice. To test this, *Pgr^cre/+^mPgrB^LsL/+^* mice exhibiting an ovarian tumor via ultrasound imaging were treated chronically for 8 weeks with placebo or the PGR antagonist, RU486. Tumor growth was measured weekly as described above. The treatment with RU486 dramatically decreased the size of the tumors compared to control (Figure 6A). After 8 weeks of treatment or until the allowable end point was reached, mice were euthanized, and tissue was harvested. Vehicle treated *Pgr^cre/+^mPgrB^LsL/+^* mice were often euthanized due to moribundity before the end of the 8-week study. However, none of the RU486 treated mice exhibited moribundity before the end of the experiment. The gross tumor morphology for *Pgr^cre/+^mPgrB^LsL/+^* animals treated with placebo or RU486 is depicted in Figure 6B-C. Ultrasound still images from the initiation of the 8-week treatment to the completion of the study demonstrate dramatic changes in size and density of the placebo and RU486 treated tumors (Figure 6D-G). ImageJ analysis of the ultrasound still images at the completion of the study reported increased cell area, cell signal, and integrative density in placebo treated tumors compared to RU486 treated tumors (Supplemental Figure 5A-C).

**Figure 6:**
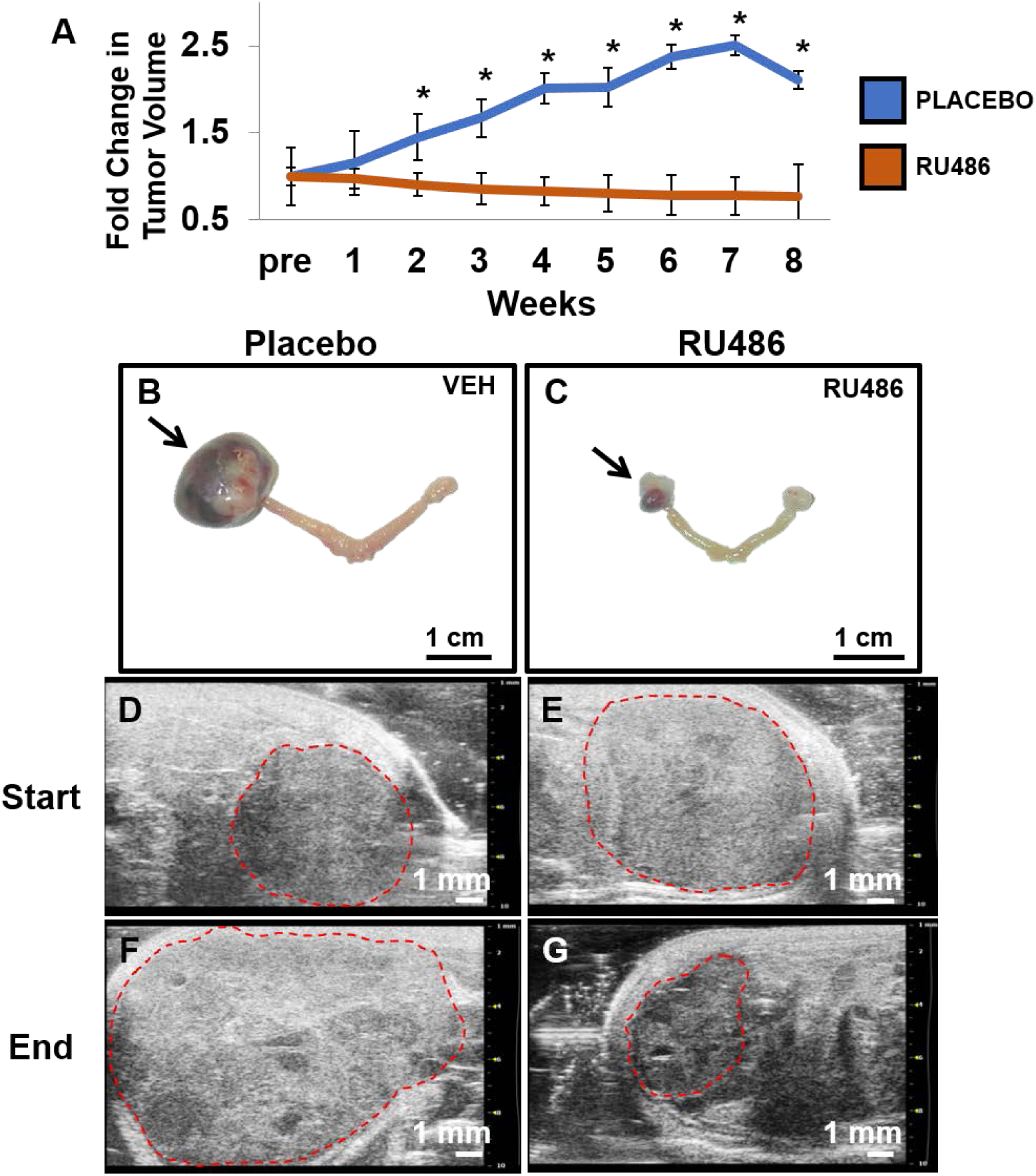
Chronic treatment of *Pgr^cre/+^mPgrB^LsL/+^* mice with RU486 abrogates tumor growth. (A) Graphical description of the fold change in tumor volume over time for *Pgr^cre/+^mPgrB^LsL/+^* tumor burdened mice treated with placebo versus the progesterone receptor antagonist, RU486. The x-axis describes the length of treatment (weeks) before sacrifice. “Pre” denotes the time before start of treatment. (B-C) Gross morphological images of placebo treated (B) and RU486 treated (C) uterus with right ovaries and left ovarian tumors. Arrows indicate the ovarian tumors. (D-G) Ultrasound images of *Pgr^cre/+^mPgrB^LsL/+^* ovarian tumors with demarcated tumor boundaries before the start of treatment (“Start”) (D-E) and after 8 weeks of treatment (“End”) (F-G) for placebo treated (D,F) versus RU486 treated tumors (E,G). The white signal observed on the ultrasound image is indicative of solid tissue, while dark regions indicate fluid-filled or less dense regions. *=denotes significance with a p-value≤0.05. Error bars represent ±SEM. Fold tumor volume change for placebo and RU486 treated tumors was calculated as the average fold change (log10(tumor volume) normalized to one) for each treatment. N=6

### Acute RU486 treatment sufficiently reduces proliferation in *Pgr^cre/+^mPgrB^LsL/+^* mouse ovarian tumors

*Pgr^cre/+^mPgrB^LsL/+^* mice exhibiting a tumor size between 0.5-1cm in diameter were administered a subcutaneous pellet of RU486 or placebo for 24 hours. Mice were euthanized and the tumor was harvested and sliced in half for nucleic acid extraction and histological fixation. A 24-hour treatment of RU486 was sufficient to attenuate the expression of the PGR target gene, *Hand2* (Figure 7A). Gross tumor morphologies for *Pgr^cre/+^mPgrB^LsL/+^* mice treated with RU486 or placebo for 24 hours are displayed in Figure 7B-C. To assess the effects of inhibition of progesterone signaling on tumor growth and cell survival, cellular proliferation and cell death number were evaluated. Within 24 hours of RU486 treatment, cell proliferation measured by BrdU incorporation was significantly attenuated compared to placebo (Figure 7D-F). Furthermore, TUNEL analysis reported a higher level of fractionated DNA fragments in the RU486 treated group compared to the placebo group (Figure 7G-K). Therefore, a 24-hour treatment of RU486 is sufficient to reduce cellular proliferation and increase cell death in tumors from *Pgr^cre/+^mPgrB^LsL/+^* mice.

**Figure 7:**
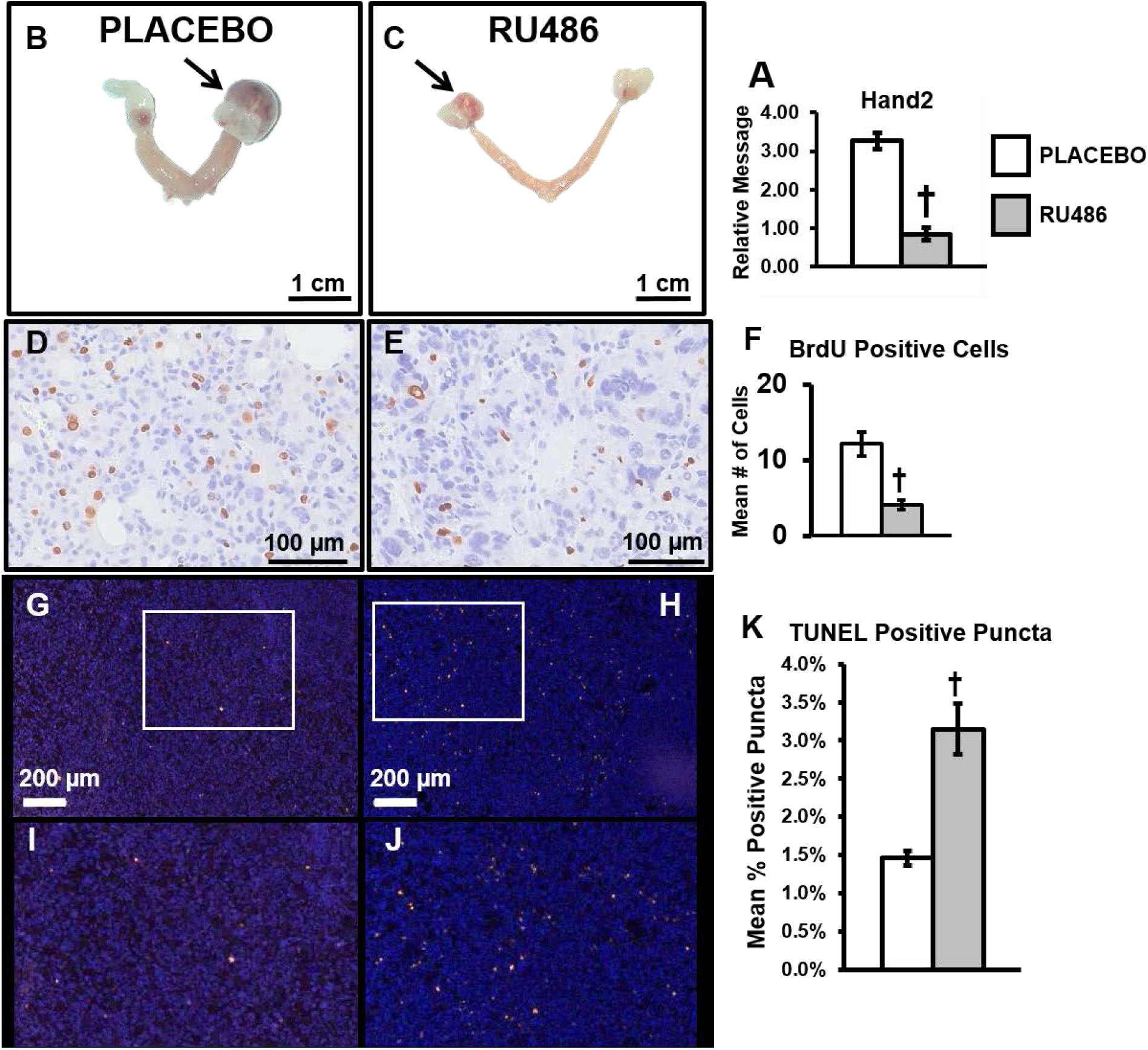
Acute RU486 treatment sufficiently prevents cell proliferation and induces cell death in *Pgr^cre/+^mPgrB^LsL/+^* tumor tissue. (A) Relative mRNA level for PGR target, *Hand2*. (B-C) Gross morphological images of vehicle (B) and RU486 (C) 24-hour treated uteri exhibiting ovarian tumors. Ovarian tumors indicated by arrows. (D-E) BrdU incorporation in tumor tissue from vehicle (D) and RU486 (E) treated tumors. (F) Quantification of BrdU positive cells from vehicle versus RU486 treated *Pgr^cre/+^mPgrB^LsL/+^* tumor tissue. (G-H) TUNEL staining from vehicle (G) and RU486 (H) 24 hour treated *Pgr^cre/+^mPgrB^LsL/+^* tumor tissue. (I-J) inset of TUNEL staining from (G-H). (K) Quantification of positive TUNEL staining in vehicle versus RU486 24 hour treated *Pgr^cre/+^mPgrB^LsL/+^* tumors. † denotes significance with a p-value ≤ 0.05. Error bars represent ±SEM. N=3

### PGRB transcriptionally regulates genes involved in cell cycle, DNA recombination, and cancer

To further understand how progesterone signaling drives tumor progression within the ovary, RNA was isolated from placebo and RU486 treated *Pgr^cre/+^mPgrB^LsL/+^* ovarian tumors and a microarray was performed. The number of differentially regulated genes totaled 995 with 383 and 612 genes upregulated and downregulated, respectively, with fold changes above 1.14 or below 0.714 with a p-value of ≤ 0.05 (Figure 8A, red circle). The list of differentially regulated genes is reported in the supplemental Excel spreadsheet, workbook 1.4. Ingenuity Pathway Analysis demonstrated the top regulated pathways included cell cycle related pathways along with breast and pancreatic cancer signaling (Supplemental Table 2). This focused list of oncogenic pathways provides a robust indication of the tumorigenic agenda of the progesterone-driven neoplasia arising in the *Pgr^cre/+^mPgrB^LsL/+^* mice.

**Figure 8:**
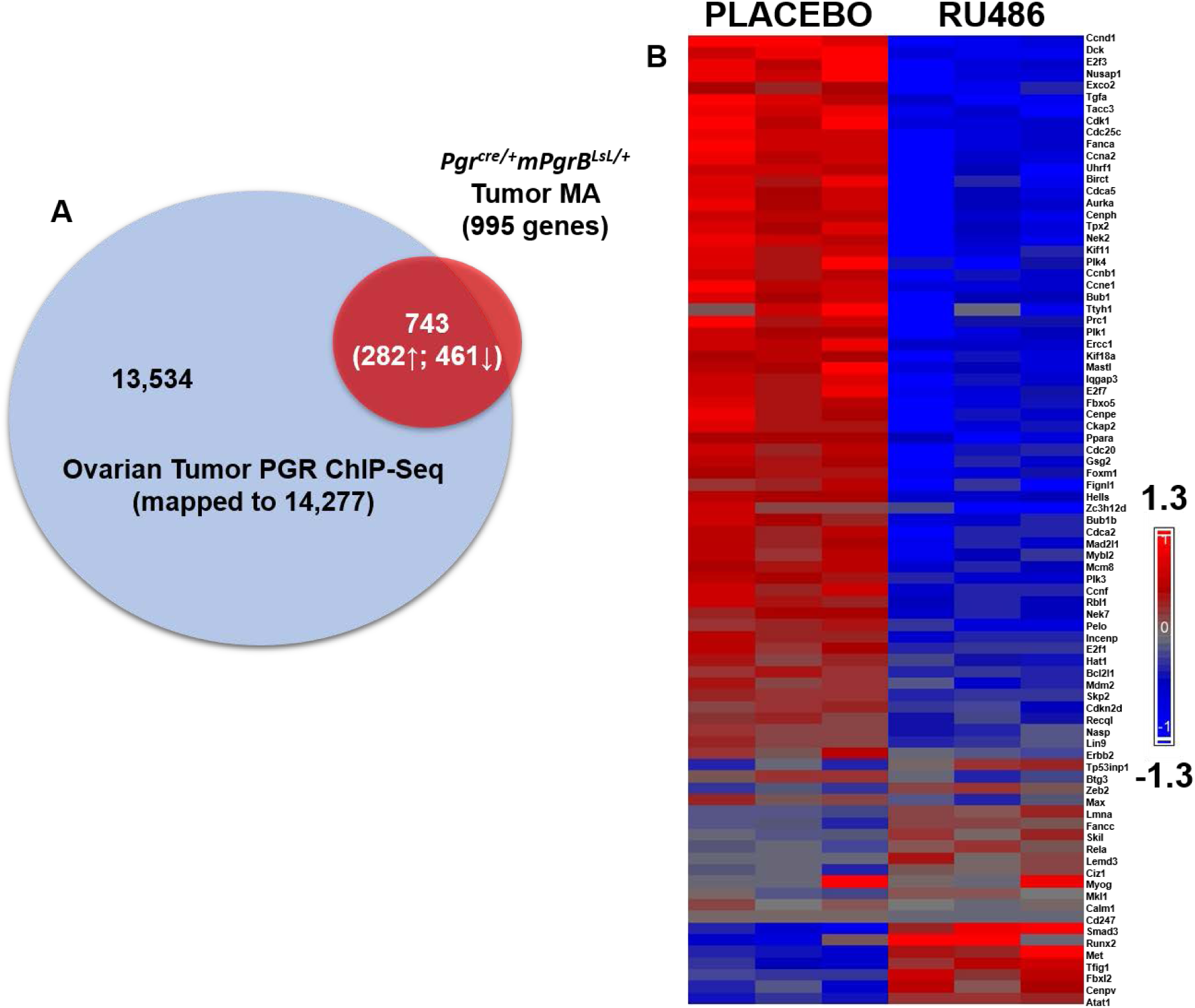
Transcriptomic and cistromic analyses define a subset of genes differentially regulated and directly bound by PGRB in the *Pgr^cre/+^mPgrB^LsL/+^* ovarian neoplasia tissue. (A) PGR ChIP-Seq data is represented by the blue circle describing the total genes bound by PGR in the enhanced promoter region of the ovarian tumor (14,277 total genes). The red circle signifies RNA microarray data displaying differentially regulated gene transcripts in the vehicle versus RU486 tumor tissue with fold changes above 1.14 or below 0.714 with a p-value of ≤ 0.05 (995 total genes). The overlapping region of both circles describes the differentially regulated genes observed in the RNA microarray and the positively bound genes identified in the ChIP-Seq (743 genes total, 282 upregulated, 461 downregulated). (B) Heat map describing the differentially regulated cell cycle genes identified via pathway analysis. Data were centered on a median of zero and color intensities were set to 1.3 and −1.3, respectively.

In order to further confirm the direct regulation of these top pathways by PGRB, a PGR ChIP-Seq analysis was performed on *Pgr^cre/+^mPgrB^LsL/+^* tumor tissue harvested from >7.5 month old virgin mice. Using the Cistrome Analysis Pipeline, the ChIP-Seq exhibited robust binding enrichment in the promoter and 5’ UTR regions of genes (Supplemental Figure 6A), with preferential localization at nuclear receptor, ATF/JUN, GATA, and CEBP motifs (Supplemental Figure 6B).

Comparative analysis was performed using the PGR bound genes from the ovarian tumor ChIP-Seq and the differentially regulated RU486 versus placebo treated tumor microarray. Of the 995 differentially regulated genes, 743 were directly bound by PGR within 5kb of the transcription start site (Figure 8A). Ingenuity Pathway Analysis of the 743 genes revealed PGR directly regulates genes involved in cell cycle, cell development, DNA recombination, and cancer pathways (Supplemental Table 3). Due to the abundant changes in cell cycle in RU486 versus placebo treated tumor tissue, a heatmap was generated to report normalized individual tumor sample values for cell cycle related genes (Figure 8B). This heatmap depicts over 80 cell cycle associated genes, many of which exhibit downregulation upon treatment with RU486. Thus, PGRB promotes the upregulation of many genes involved in the cell cycle, a function that is attenuated upon treatment with RU486.

### PGRB directly promotes cell cycle progression through activation of CCND1

Upon acute treatment with RU486, *Pgr^cre/+^mPgrB^LsL/+^* ovarian tumors exhibited decreased transcript levels of *Ccnd1* (Figure 9A), the critical cyclin necessary for entry into S phase of the cell cycle^18^. To confirm this potential change in CCND1 protein expression, western blot analysis was performed on *Pgr^cre/+^mPgrB^LsL/+^* ovarian tumors acutely and chronically treated with RU486 or placebo. Western blot results depicted slightly attenuated levels of CCND1 upon acute treatment with RU486 (Figure 9B). However, chronic RU486 treatment resulted in a dramatic decrease of CCND1 protein levels compared to placebo (Figure 9B). Furthermore, the PGRB ChIP-Seq analysis confirms the presence of a strong PGR binding event on the murine *Ccnd1* locus (Figure 9C). ChIP-qPCR at this region 226 bp upstream of the transcription start site confirmed efficient PGR binding (Figure 9D). Additionally, Ingenuity Pathway Analysis of the tumor microarray revealed multiple downstream targets of CCND1 also exhibiting attenuation upon treatment with RU486 (Figure 9E).

**Figure 9:**
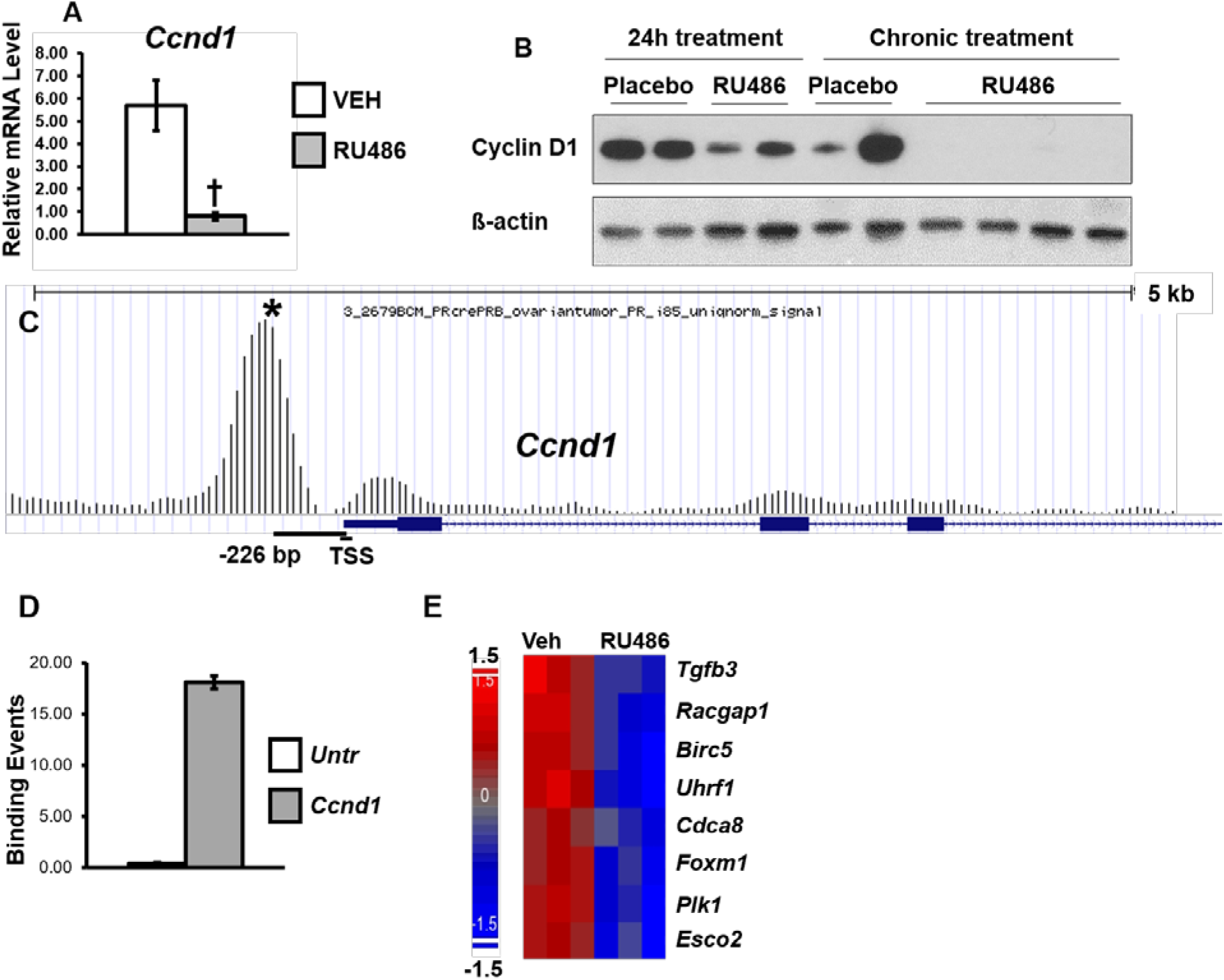
PGRB directly regulates CCND1 expression through binding to the *Ccnd1* locus. (A) Relative mRNA levels for *Ccnd1* after acute RU486 or placebo treatment of *Pgr^cre/+^mPgrB^LsL/+^* ovarian tumors. (B) Western blot analysis of *Pgr^cre/+^mPgrB^LsL/+^* tumors treated with RU486 or placebo for 24 hours or 8 weeks (chronic). Total CCND1 and β-actin protein are displayed. (C) Graphical description of the PGRB binding event on the loci of *Ccnd1*. (D) ChIP-qPCR validation of the PGRB binding event 226 bp upstream of the transcription start site (TSS) at the *Ccnd1* locus. Y-axis indicates the amount of binding events per 1000 cells in the untranslated region versus interval of interest. Error bars indicate the standard deviation of averaged binding occurrences. (E) Heat map describing the differentially regulated *Ccnd1* gene targets identified via pathway analysis. Data were centered on a median of zero and color intensities were set to 1.5 and −1.5, respectively. † denotes significance with a p-value ≤ 0.05. Error bars represent ±SEM. Asterisk indicates the binding interval validated in (D).

Of these target genes, FOXM1 and PLK1 are both critical proteins necessary for the initiation of mitosis or M phase^19, 20^. Due to their strong role in mitotic entry, both are found upregulated in many cancers^21, 22^. To identify the role of PGRB in the promotion of mitosis, RNA transcript levels of known mitotic-initiating proteins were evaluated. Acute RU486 treatment of *Pgr^cre/+^mPgrB^LsL/+^* ovarian tumors results in the attenuation of multiple genes involved in M phase entry including *Foxm1*, *Plk1*, *Cdc25c*, *Ccnb1*, and *Cdk1* (Supplemental Figure 7A-E), providing further explanation for the decreased proliferation observed in these tumors. Additionally, PGR ChIP-Seq analysis confirmed PGR binding events at the loci of *Foxm1*, *Plk1*, *Cdc25c*, and *Cdk1* (Supplemental Figure 7F-I), which were confirmed via ChIP-qPCR analysis (Supplemental Figure 7J-M). Therefore, PGRB strongly regulates CCND1 and proteins involved in the progression of mitosis, resulting in an increase of cell division.

## Discussion

Through the utilization of mice expressing high levels of PGRB, we have provided evidence for the PGRB specific promotion of a cell cycle gene signature resulting in upregulated cellular proliferation and uncontrolled growth. The PGRB isoform binds directly to the *Ccnd1* promoter and also promotes the AKT pathway, resulting in rapid transition states during M and G1 phases, causing persistent cell growth. This work describes a unique role for the PGR isoforms in the initiation and progression of ovarian neoplasia, with a molecular signature similar to that of human ovarian adenocarcinoma. Therefore, this article describes a robust, *in vivo* role for the PGRB isoform in the promotion of cellular proliferation in endocrine tissues and development of ovarian neoplasia.

Although ovarian PGR expression in wildtype mice is limited to granulosa cells of the pre-ovulatory follicle^23^, we observed expression of the PGRB knock-in allele in corpora lutea, clearly observed in ovaries from *Pgr^cre/+^Rosa^mT/mg^* mice. Activated expression of PGRB by Cre recombinase in the follicle likely maintained expression in the corpora lutea post differentiation^24^, as PGR is not normally expressed in the corpora lutea of the rodent^25, 26^. The unexpected expression pattern of PGR in the ovary was likely a foreshadowing of future events, as mice developed abnormal cells at the pre-neoplastic stage that progressed to tumors at 23 weeks of age in the neoplastic stage. These abnormal growths were not only positive for PGR, but also exhibited high levels of the proliferative marker, Ki67, with a decreased amount of the apoptotic marker, CASP3. Therefore, appropriate expression levels of the PGRB isoform are important for maintenance of normal ovarian tissue.

The majority of ovarian cancers in humans are derived from the epithelium and are classified into five different carcinoma subtypes^27^, yet the tumors observed within the *Pgr^cre/+^mPgrB^LsL/+^* mice appear to arise from granulosa cells that might have failed to undergo terminal differentiation during the process of luteinization. Tumors originating from granulosa cells, such as granulosa cell tumors and rare luteomas, can develop from genetic predisposition and aberrant growth factor and hormone levels^28^. Furthermore, the ovarian neoplasms from the *Pgr^cre/+^mPgrB^LsL/+^* mice are sensitive to the PGR antagonist, RU486, confirming that these tumors require progesterone signaling for growth. Interestingly, the tumors identified in the *Pgr^cre/+^mPgrB^LsL/+^* mice expressed high levels of AKT signaling with a gene expression signature most consistent with human ovarian adenocarcinoma, according to the GSEA and NextBio comparative analyses. Accordingly, high AKT levels may be a result of impaired ARID1A levels, a member of the SWI/SNF family with a proposed role in protecting against progesterone resistance^29^.

The PGR isoforms have consistently exhibited different functions within the endocrine organs^30^. Notably, PGRB has often been associated with proliferation in the mammary gland while PGRA exhibits anti-proliferative and anti-inflammatory functions in the uterus. Due to the proliferative functions of PGRB, it is not unexpected to observe progesterone-driven ovarian tumors in the *Pgr^cre/+^mPgrB^LsL/+^* mice. However, the *Pgr^cre/+^mPgrA^LsL/+^* mice also develop abnormal cells and ovarian tumors at 33 weeks of age (Table 1), albeit at a much lower frequency and independent of abnormal progesterone, FSH, and LH levels (Supplemental Figure 4). Thus, it is important to understand how sustained PGRA or PGRB expression in the ovarian corpora lutea can elicit ovarian neoplasia. In rodents, the corpus luteum is maintained throughout the duration of pregnancy and produces high levels of progesterone hormone and a small amount of estrogen to preserve the pregnancy^31^. The presence of this progesterone promotes its own production^26^ and simultaneously inhibits cellular apoptosis within corpora lutea^32^, making it a pro-survival factor. Indeed, progesterone in the corpora lutea was identified to dose-dependently prevent Fas-mediated apoptosis in corpus luteum regression^33^. Regression of corpora lutea occurs first by a decrease in active progesterone and then an activation of the Fas pathway, resulting in cellular apoptosis. Thus, active progesterone signaling is responsible for maintaining the life of the corpus luteum. In normal states, nuclear PGR is not expressed in normal corpora lutea, yet progesterone may be signaling via the membrane PGR family in rodents^34^. However, in the *Pgr^cre/+^mPgrB^LsL/+^* mice, with the addition of nuclear PGR in the corpora lutea, luteal regression may be delayed due to the progesterone-driven survival mechanism. Although virgin *Pgr^cre/+^mPgrB^LsL/+^* mice only maintain corpora lutea for two days each estrous cycle^31^, the presence of nuclear PGR and large amounts of progesterone ligand after 20 weeks of age may create the suitable environment for uncontrolled cellular growth in the ovarian stroma. This hypothesis also correlates with the observed decrease in the apoptotic marker, CASP3, and increased expression of proliferation markers in advanced stages in the ovary from *Pgr^cre/+^mPgrB^LsL/+^* mice (Figure 2). Therefore, further work is required to investigate whether expression of nuclear PGR in the corpora lutea causes delayed luteal regression and maintained cellular growth.

Utilization of selective hormone receptor modulators can be beneficial in the treatment of ovarian neoplasms (reviewed in^2, 35^). RU486, a progesterone receptor inhibitor, has demonstrated effectiveness in minimizing drug resistant ovarian cancer^36^. Furthermore, RU486 has also been tested in the treatment of breast and endometrial cancers (reviewed in^2^). Through utilization of a chronic RU486 treatment, the *Pgr^cre/+^mPgrB^LsL/+^* tumors experienced shrinkage over an 8-week time period. In addition to reduced growth, RU486 treated tumors exhibited a strikingly less dense appearance using ultrasound imaging compared to the placebo group. The placebo treated animals were consistently found moribund or dead before the treatment endpoint, providing further evidence that RU486 was successful in mitigating tumor growth and progression of disease in these animals. An acute treatment with RU486 was also successful in the stalling of tumor growth in the *Pgr^cre/+^mPgrB^LsL/+^* mice. Cells within RU486 treated tumors exhibited a lower proliferation index and heightened levels of cellular death by DNA fragmentation markers. The RNA microarray of RU486 and placebo treated *Pgr^cre/+^mPgrB^LsL/+^* tumors demonstrated a total of 995 genes differentially regulated, with 743 genes exhibiting direct transcriptional regulation by the PGR. Within just a 24-hour RU486 treatment, multiple pathways were found dysregulated, including an assortment of about 80 cell cycle related genes favoring the slowing or stopping of the cell cycle. Thus, PGRB signaling in the ovary is a potent regulator of the cell cycle pathway.

The PGRB has previously exhibited a role for increased proliferation in the uterus^8^ and in the mammary gland^7^. The regulation of the S-phase entry cyclin, *Ccnd1*, by the PGR has been exhaustively investigated using *in vitro* immortalized breast cancer cells^37–41^ and has been confirmed in murine mammary epithelial tissue^42^. However, positive correlation between PGR and CCND1 expression patterns has been variable and sometimes contradictory in human breast carcinoma^43–45^. Our data provide a clear role for efficient binding of PGRB on the murine *Ccnd1* promoter region. Furthermore, treatment with RU486 results in decreased CCND1 protein expression over time, suggesting PGRB is prevented from promoting transcription of the *Ccnd1* locus. Therefore, PGRB directly regulates the progression of the cell cycle through transcriptional activation of *Ccnd1*.

Additionally, CCND1 downstream targets, *Plk1* and *Foxm1*, also exhibited attenuation upon acute treatment with RU486. Active PLK1 plays an integral role in cell division through preparing the cell for mitosis by phosphorylating CDC25C, which subsequently activates the CDK1-CCNB1 complex^20, 46, 47^. PLK1 is also indispensable for spindle assembly chromosome separation, progression of anaphase, and cellular cytokinesis. Previous data have described an additional role for FOXM1 in the activation of PLK1 for mitotic entry, through the identification of forkhead binding sites on the PLK1 promoter^48^. Indeed, it was concluded that FOXM1 not only regulates PLK1, but both are involved in a positive feedback loop to promote mitosis^19^. This work provided further detailed analysis of PGR binding directly at the loci of mitosis-promoting genes: *Foxm1*, *Plk1*, *Cdc25c*, and *Cdk1*. Since PLK1 and FOXM1 are necessary for cell cycle progression^19^, they have consistently been identified to be expressed in a variety of cancers^21, 22^. PLK1 alone is found upregulated in human ovarian cancer^49, 50^, endometrial carcinoma^51^, ectopic endometriosis^52^, and breast cancer^53^, often correlating with increased proliferation and severity. Further work is required to understand the function of PGRB-driven epithelial proliferation within the uterus and whether PLK1 mediates this unique function of PGRB in the normal reproductive tract. Therefore, the regulation of mitosis by the PGRB isoform provides an interesting mechanism to explain the growth of ovarian neoplasia and additional non-invasive, PGR-positive, solid tumors arising in endocrine organs.

Within this study, we have generated a unique PGRB expressing mouse model exhibiting the development of ovarian neoplasms at 23 weeks of age. In these mice, PGRB promoted cellular proliferation through the control of many cell cycle regulatory genes in the ovarian environment. Specifically, S phase initiator, CCND1, and critical regulators of the G2/M transition, FOXM1, PLK1, CDC25C, CCNB1, and CDK1, were all evidenced to be strongly regulated by PGRB expression (model depicted in Figure 10). Additionally, these progesterone-driven ovarian growths exhibited a strong correlative gene expression signature with human adenocarcinoma driven by aberrant AKT signaling. AKT signaling is a potent promoter of S-phase entry and has the potential to be regulated by a non-canonical method of PGR signaling^54^ (Figure 10). In conclusion, these data describe a novel mechanism for the direct regulation of cellular proliferation by PGRB, providing valuable insight into the uncontrolled growth of PGRB-positive neoplasia occurring in women today.

**Figure 10:**
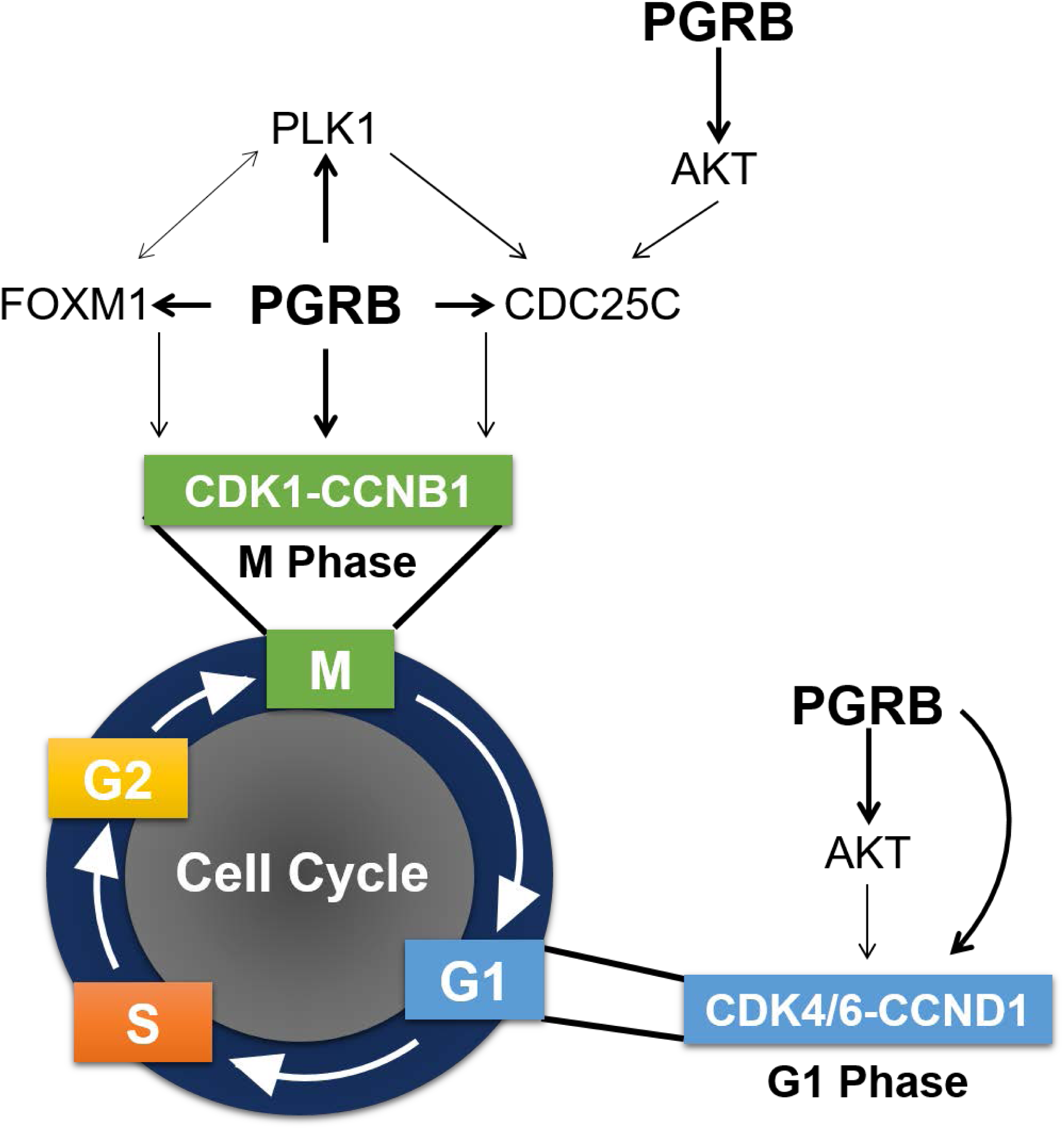
PGRB promotes growth through regulation of the cell cycle pathway. Graphical model portraying the role of PGRB in the initiation of M and G1 phases via regulation of multiple genes responsible for mitotic entry.

## Supporting information

DEG List

## Acknowledgement

The authors thank Sungnam Cho, Jie Yang, Yiqun Zhang, Elise Mangin, Arturo Garza, and Janet DeMayo for technical assistance. This work was impossible without the support of the Core resources at the NIEHS and Baylor College of Medicine. These include the Integrative Bioinformatic Support Group, Knockout Mouse Core, Comparative Medicine Branch Epigenomic and DNA Sequencing Core, Molecular Genomic Core, Clinical Pathology Group and the Pathology Support Group at the NIEHS Also the Embryonic Stem Cell Core, the Genetically Engineered Mouse Core, Mouse Phenotyping Core, and the Genomic and RNA Profiling Core at Baylor College of Medicine. This work was supported by NIH Grants: Z1AES103311-01, R01HD042311, CA125123 (to CJC), 5U54HD007495 (to FJD) and R01HD042311 (to JPL) and NURSA grant: U19DK62434 (to FJD) and CIHR grant 125936 (to BDM).

---

**Supplemental Figure 1:**
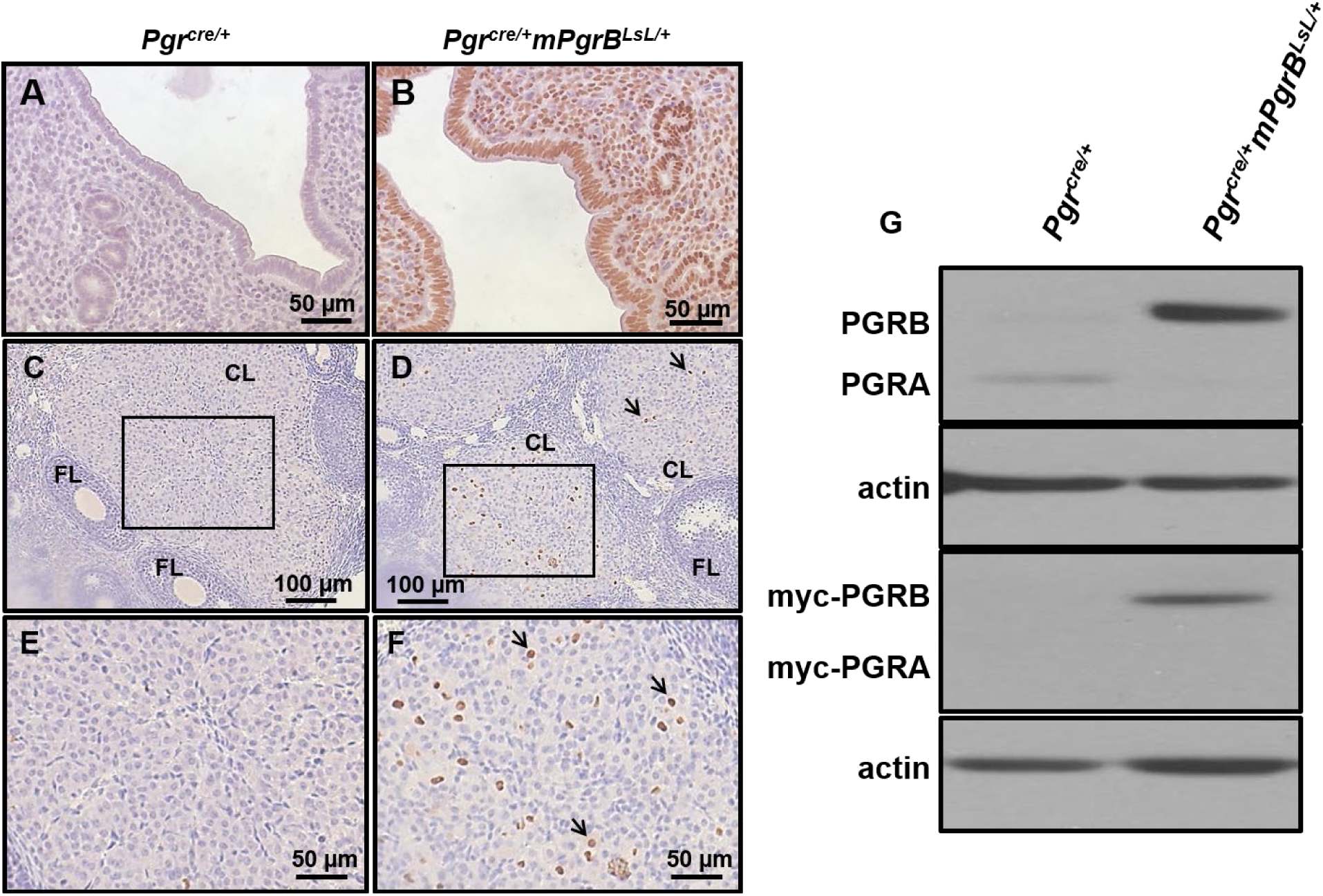
Expression levels of the conditional overexpression allele for *Pgr*B in uterine and ovarian tissue. (A-D) Myc-tag immunohistochemistry of 8 week old wildtype uterus (A) and ovary (C) compared to *Pgr^cre/+^mPgrB^LsL/+^* mouse uterus (B) and ovary (D). (E-F) High magnification images of corpora lutea from insets in (C-D) in wildtype ovary (E) and *Pgr^cre/+^mPgrB^LsL/+^* ovary (F). (G) Western blot depicting PGR and myc-tag protein levels from whole uterine isolates in wildtype *Pgr^cre/+^* and *Pgr^cre/+^mPgrB^LsL/+^* mice. PGRB protein (118 kDa), PGRA protein (90 kDa)

**Supplemental Figure 2:**
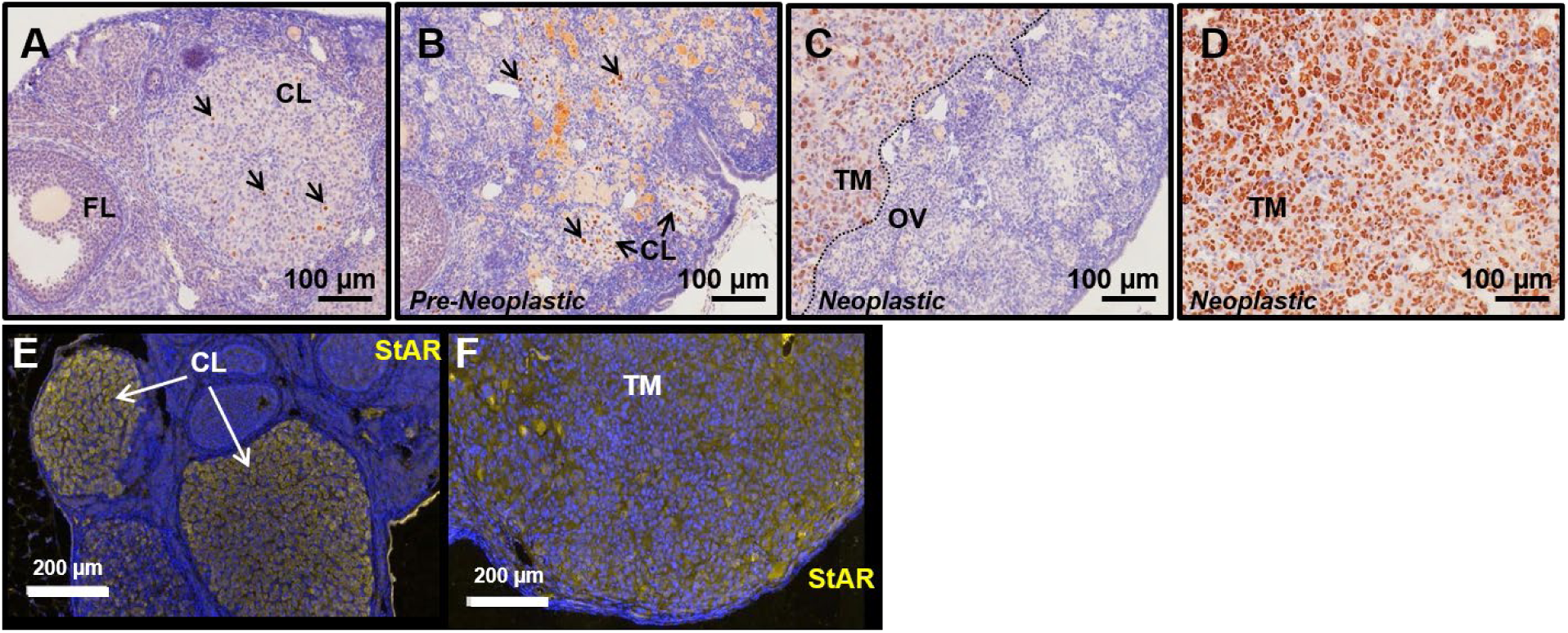
PGR positive ovarian neoplasia from *Pgr^cre/+^mPgrB^LsL/+^* mice consume the entire ovarian bursa and exhibit robust StAR expression levels. Immunohistochemical images of PGR positive cells within corpora lutea (indicated by arrows) from normal *Pgr^cre/+^mPgrB^LsL/+^* mouse ovaries at 10 weeks (A) and from pre-neoplastic ovaries (B). (C-D) PGR immunohistochemistry of PGR positive outgrowth of cells occurring in neoplastic ovaries from *Pgr^cre/+^mPgrB^LsL/+^* mice. Immunofluorescent staining for steroidogenic acute regulatory protein (StAR) in a wildtype ovary (E) and neoplastic ovarian tumor (F) of *Pgr^cre/+^mPgrB^LsL/+^* mice. Blue staining is DAPI, yellow staining represents positive StAR staining. FL=follicle, CL=corpora lutea, OV=ovary, TM=tumor tissue

**Supplemental Figure 3:**
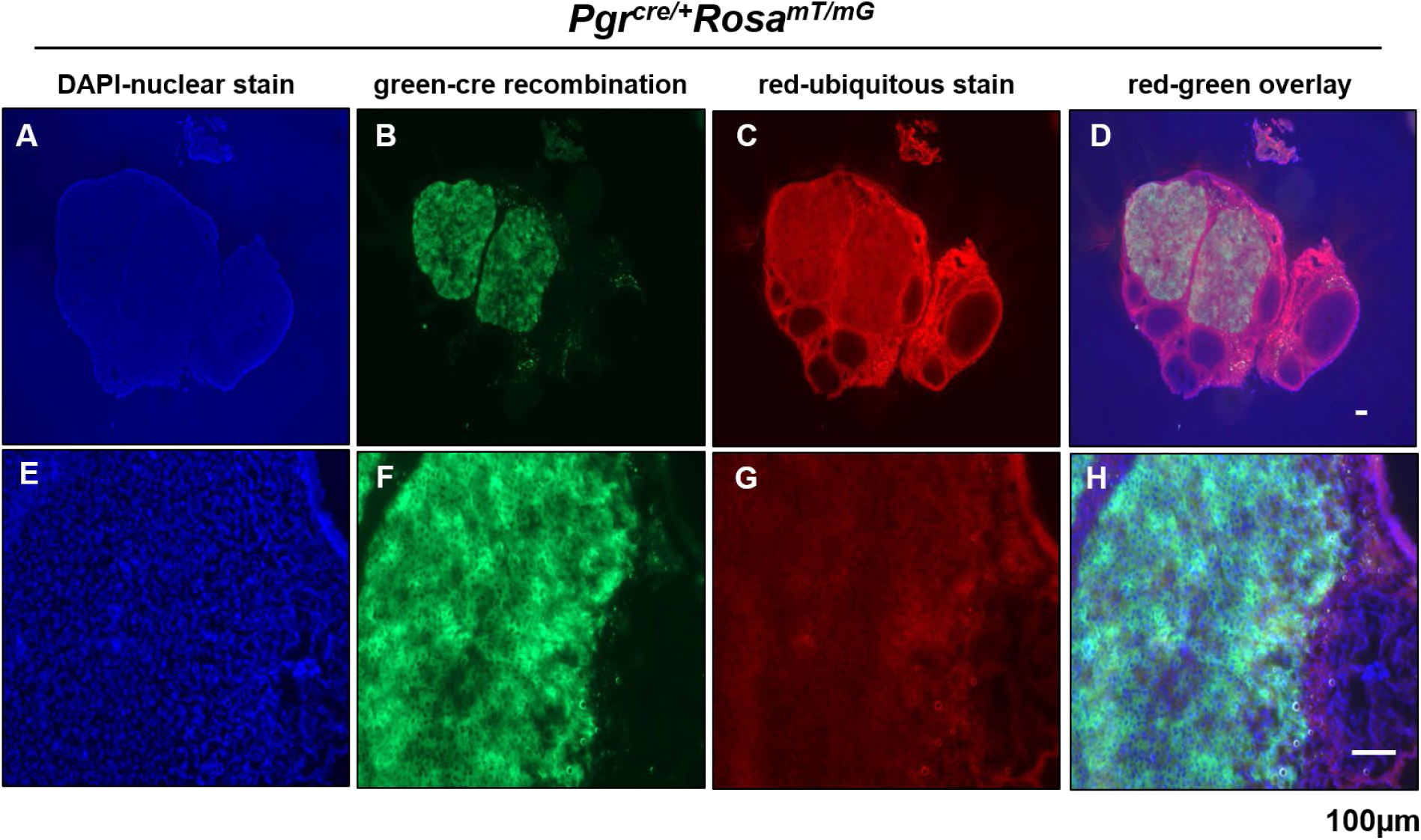
Lineage tracing using the *Pgr^cre/+^Rosa^mT/mG^* model demonstrates localized Cre recombination to the corpus luteum at 8 weeks of age. DAPI nuclear staining (A,E), positive Cre recombinase activity in green from the *Rosa^mT/mG^* construct (B,F), ubiquitous red fluorescence from *Rosa^mT/mG^* model (C,G), and red-green overlay (D,H). *Pgr^cre/+^Rosa^mT/mG^* mouse ovary (A,B,C,D) and ovarian corpus luteum (E,F,G,H).

**Supplemental Figure 4:**
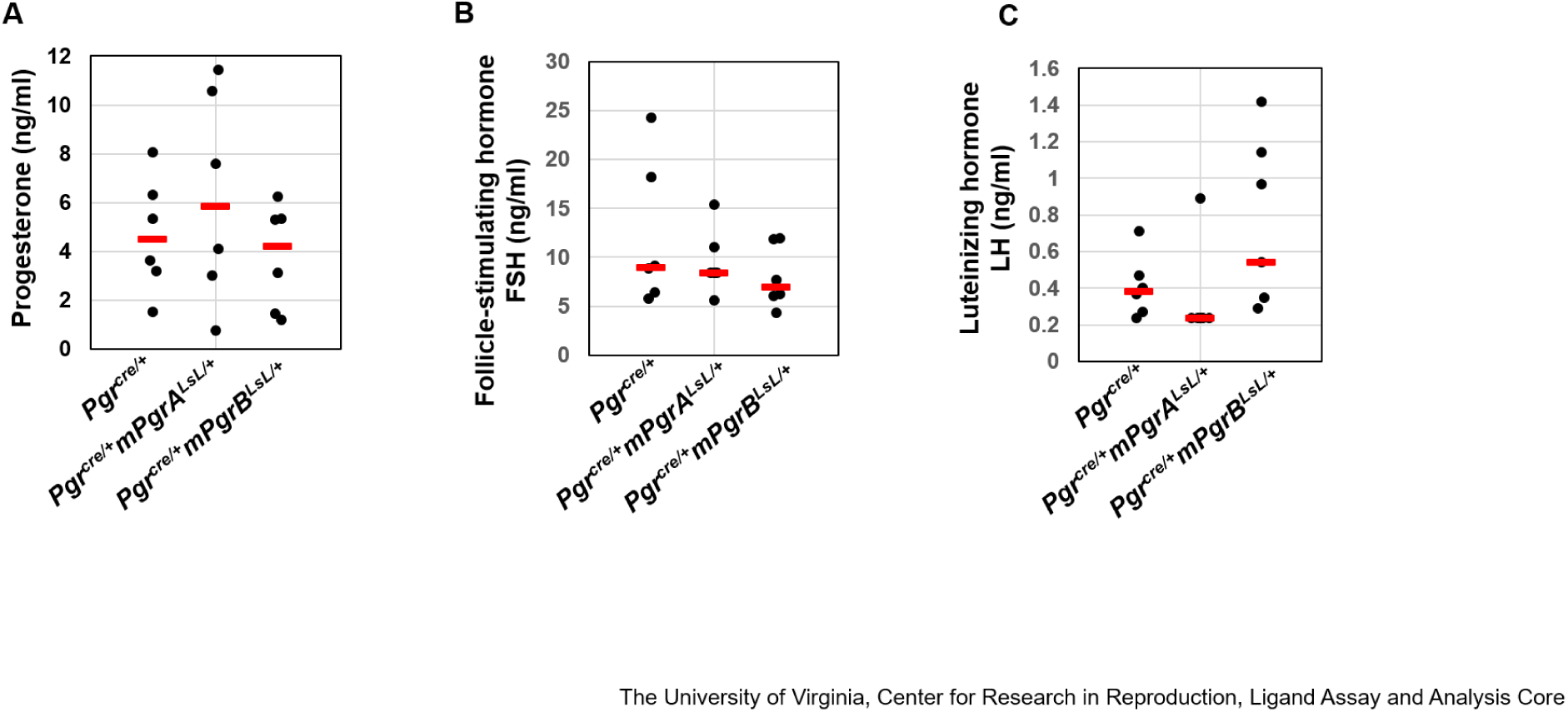
Endogenous Hormone Level Measurements. Serum levels of progesterone (A), follicle-stimulating hormone (FSH) (B), and luteinizing hormone (LH) (C) were not altered in 23 week-old *Pgr^cre/+^mPgrA^LsL/+^* mice and *Pgr^cre/+^mPgrB^LsL/+^* mice at diestrous stage. Each dot indicates the serum level from one mouse. The red line indicates median level.

**Supplemental Figure 5:**
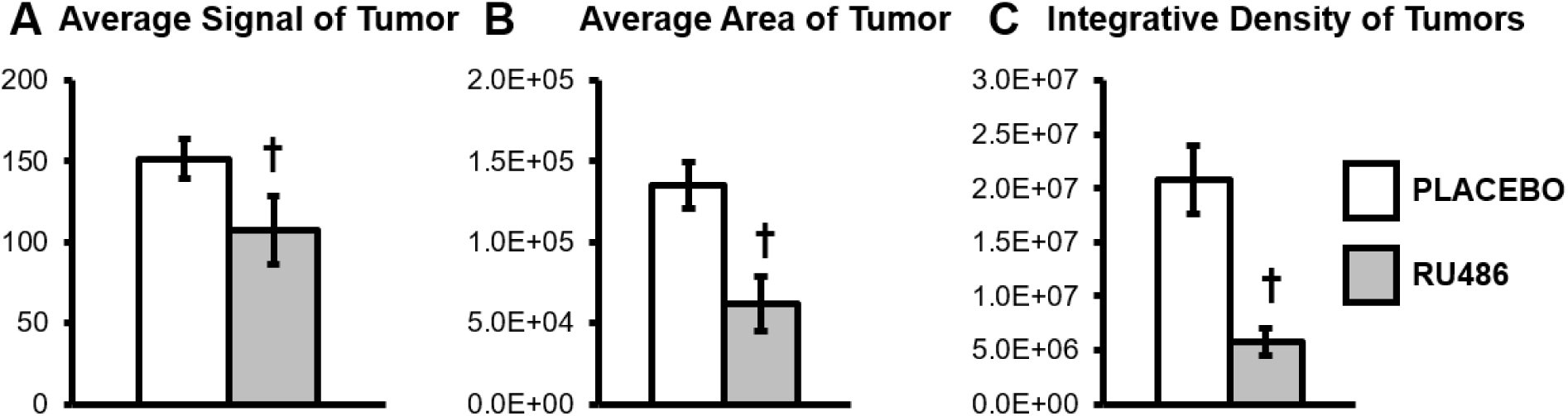
*Pgr^cre/+^mPgrB^LsL/+^* tumors exhibit decreased density and size after acute RU486 treatment. (A-C) ImageJ analysis on tumor frames exhibiting the greatest tumor diameter taken from the final week of treatment. (A) Reports average signal output or lightened areas in tumor images in RU486 vs placebo groups. (B) Depicts average areas of tumor as a function of the measurement tool used in the ImageJ software platform of RU486 versus placebo treated tumors. (C) Average integrative density (cell signal multiplied by cell area) for the ultrasound frames from RU486 versus placebo treated *Pgr^cre/+^mPgrB^LsL/+^* ovarian tumors. † denotes significance with a p-value≤0.05. Error bars represent ±SEM.

**Supplemental Figure 6:**
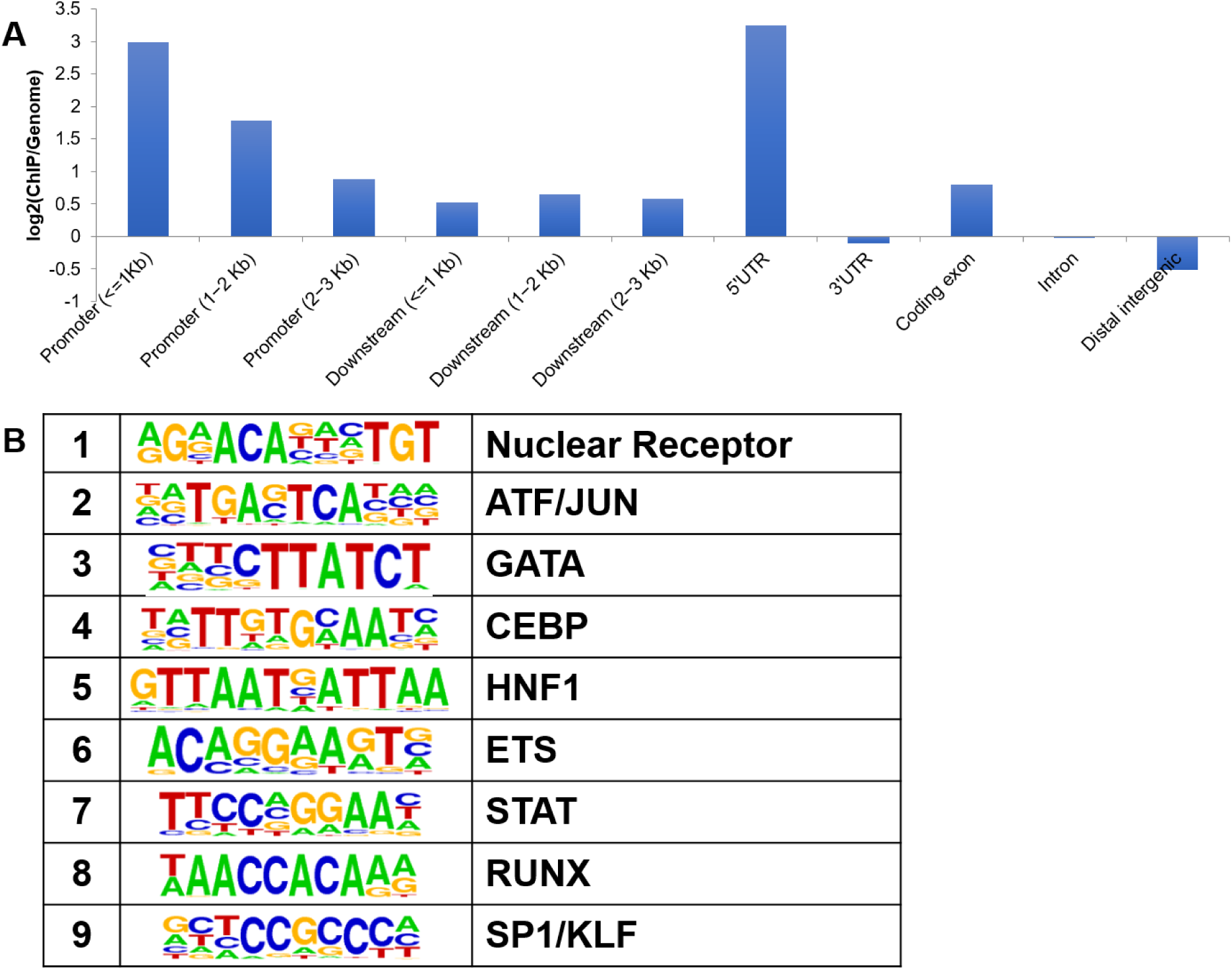
PGR ChIP-seq is enriched at promoter regions and binds to hormone receptor binding sequences. (A) Enrichment distribution of PGR binding on the genome compared to normal expected enrichment. Data is equivalent to the natural log of PGR chip binding/basal genome binding. (B) Top binding of known motifs by PGR in the ovarian tumor ChIP-Seq.

**Supplemental Figure 7:**
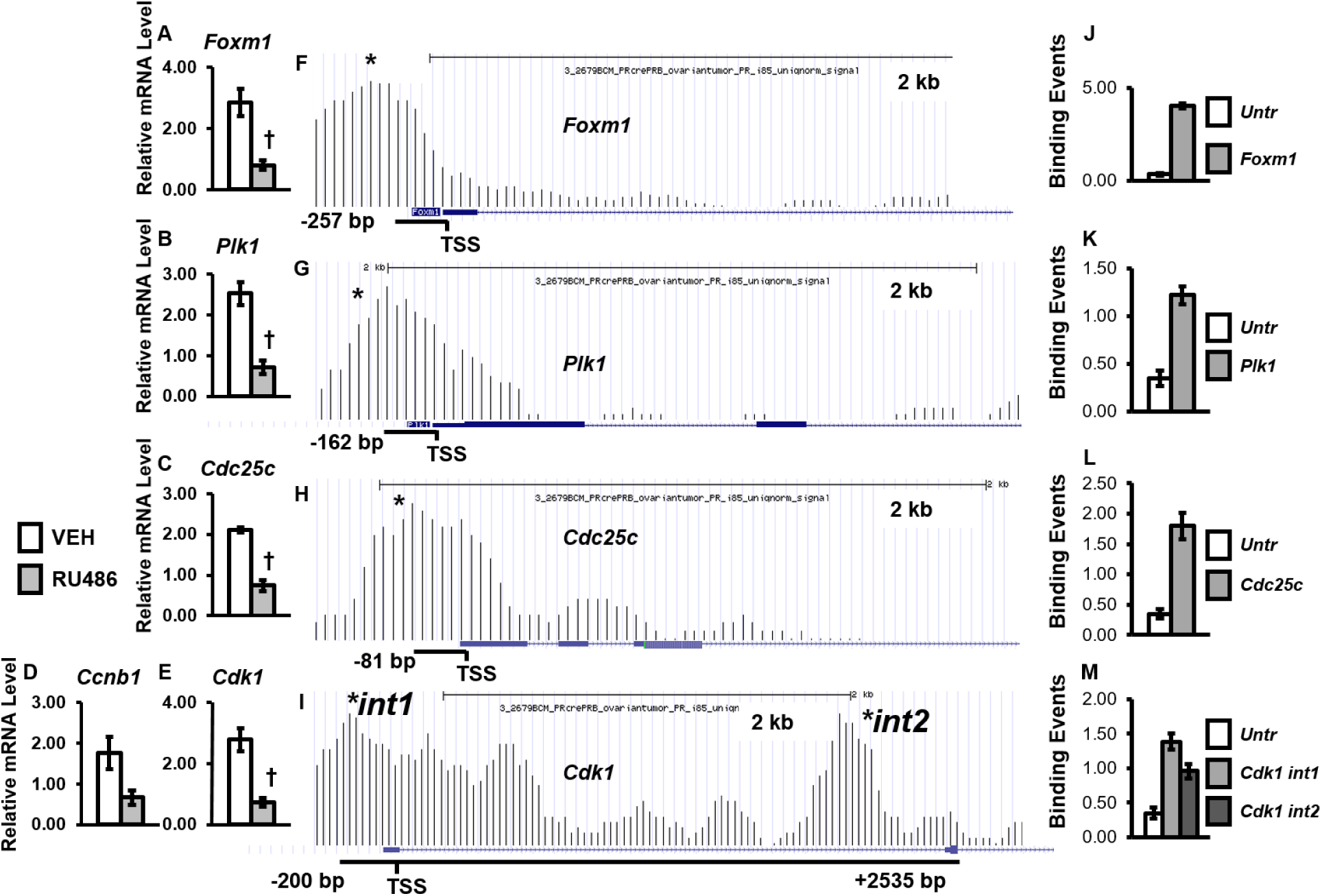
PGRB promotes the cell cycle through direct regulation of genes necessary for the G2/M transition. Relative message levels for (A) *Foxm1*, (B) *Plk1*, (C) *Cdc25c*, (D) *Ccnb1*, and (E) *Cdk1*. (F-I) Graphical description of the PGR binding events for (F) *Foxm1*, (G) *Plk1*, (H) *Cdc25c*, and (I) *Cdk1* with validated binding intervals indicated by asterisks and distances reported from transcription start site (TSS). (J-M) ChIP-qPCR validation results of binding events indicated with asterisks described in (F-I) for *Foxm1* (J), *Plk1* (K), *Cdc25c* (L), and *Cdk1* (M). Y-axis represents the number of binding events detected per 1000 cells in the untranslated region versus interval of interest. Error bars indicate the standard deviation of averaged binding occurrences.

**Supplemental Table 1:**
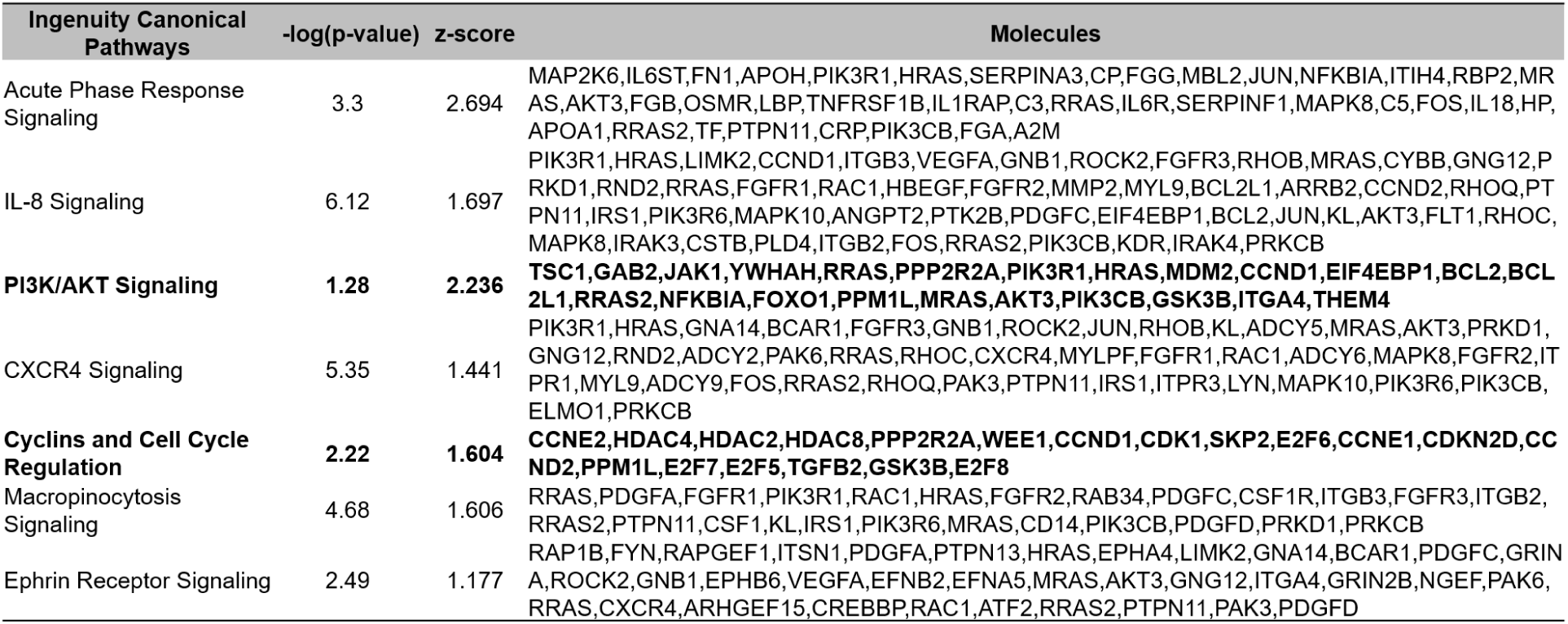
Top Altered Canonical Pathways in 23 Week Old Neoplastic *Pgr^cre/+^mPgrB^LsL/+^* Mouse Ovaries. Ingenuity Pathway Analysis was utilized to obtain the list of top canonical pathways from the RNA microarray performed on 23 week old, neoplastic *Pgr^cre/+^mPgrB^LsL/+^* mouse ovarian tumor tissue.

**Supplemental Table 2:**
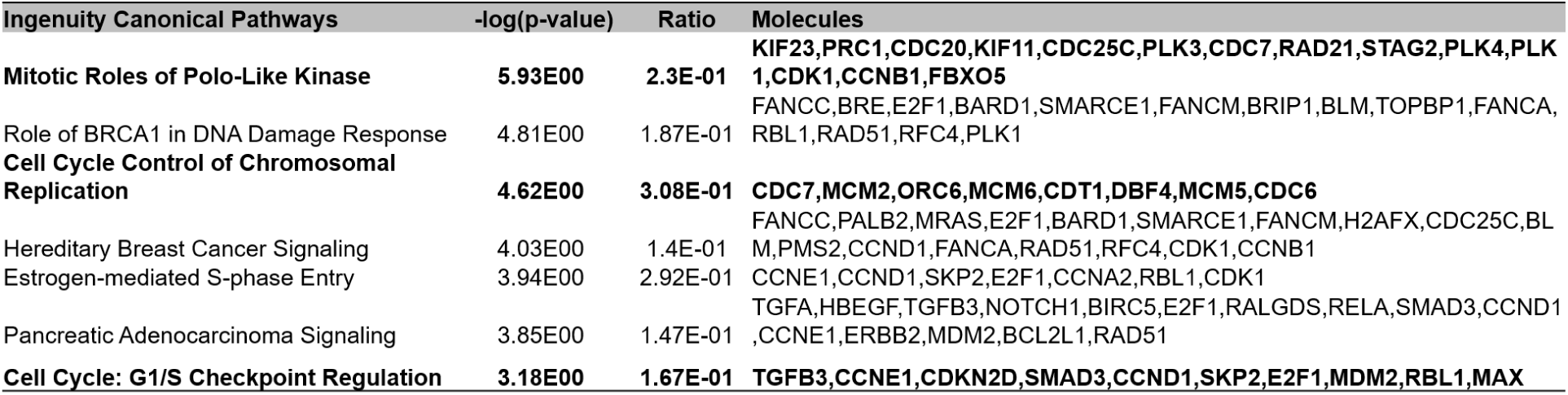
PGR regulates multiple canonical pathways involved in cell proliferation and cancer. Top canonical pathway list for differentially regulated genes between placebo and RU486 acute treatment in the RNA microarray.

**Supplemental Table 3:** PGR transcriptionally regulates multiple canonical pathways involved in cell proliferation and cancer. Top biological function gene list from Ingenuity Pathway Analysis for the tumor microarray differentially regulated gene list overlapped with the PGR ovarian tumor ChIP-seq analysis. #Mols=number of molecules, p-Value=p-value based on the Ingenuity Pathway Analysis algorithm for analyzing enrichment in a pathway.

| Categories | p-Value | # Mols |
| --- | --- | --- |
| Embryonic Development, Organismal Survival | 1.35E-10 | 28 |
| Cell Cycle | 8.44E-08 | 19 |
| Cell Cycle, DNA Replication, Recombination, and Repair | 3.67E-07 | 15 |
| DNA Replication, Recombination, and Repair | 6.22E-07 | 17 |
| Cell Cycle, Cellular Assembly and Organization, DNA Replication, Recombination, and Repair | 7.47E-07 | 7 |
| Cancer, Organismal Injury and Abnormalities, Respiratory Disease | 2.55E-06 | 27 |
| Cell Death and Survival | 2.69E-06 | 120 |
| Cellular Assembly and Organization | 3.33E-06 | 7 |
| Cellular Compromise, DNA Replication, Recombination, and Repair | 6.79E-06 | 10 |
| Cancer, Organismal Injury and Abnormalities | 6.86E-06 | 48 |

**Supplemental Table 4:**
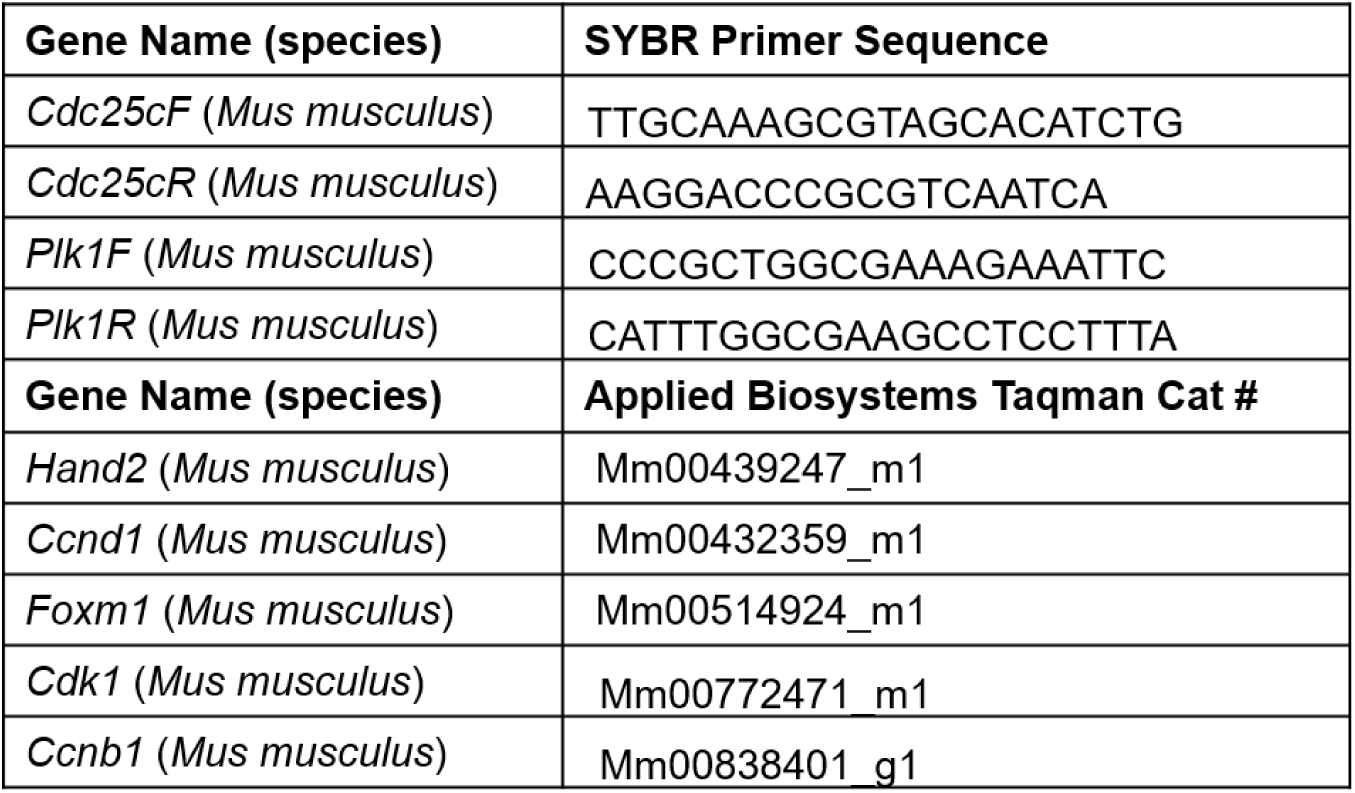
Complete list of SYBR primer sequences and Applied Biosystems Taqman probe catalog numbers.

